# DCARS: Differential correlation across ranked samples

**DOI:** 10.1101/303735

**Authors:** Shila Ghazanfar, Dario Strbenac, John T. Ormerod, Jean Y. H. Yang, Ellis Patrick

## Abstract

Genes act as a system and not in isolation. Thus, it is important to consider coordinated changes of gene expression rather than single genes when investigating biological phenomena such as the aetiology of cancer. We have developed an approach for quantifying how changes in the association between pairs of genes may inform patient prognosis called Differential Correlation across Ranked Samples (DCARS). Modelling gene correlation across a continuous sample ranking does not require the classification of patients into ‘good’ or ‘poor’ prognosis groups and can identify differences in gene correlation across early, mid or late stages of survival outcome. When we evaluated DCARS against the typical Fisher Z-transformation test for differential correlation, as well as a typical approach testing for interaction within a linear model, on real TCGA data, DCARS significantly ranked gene pairs containing known cancer genes more highly across a number of cancers. Similar results are found with our simulation study. DCARS was applied to 13 cancers datasets in TCGA, revealing a number of distinct relationships for which survival ranking was found to be associated with a change in correlation between genes. Furthermore, we demonstrated that DCARS can be used in conjunction with network analysis techniques to extract biological meaning from multilayered and complex data.

Availability: https://github.com/shazanfar/DCARS.

## INTRODUCTION

Correlation quantifies the degree of monotonic association between two variables. In the context of gene expression, correlated genes are often associated with sets of genes that are co-ordinately expressed and are likely to belong to biologically meaningful groups such as pathways or perform a similar function. In cancer, coordination of gene expression can be disrupted due to various genetic anomalies such as point mutations, copy number variations or larger structural changes in the genome (Sevimoglu and Arga, 2014; Ciriello *et al.*, 2013; Bashashati *et al.*, 2012). The level of concordance among genes in expression profiles of cancerous tissue relative to a normal tissue is one lens through which cancer can be investigated. Genes that develop a concordant pattern of expression in tumours may work together to evade anti-tumour mechanisms, for example increase in metabolic activity in tumour tissues (Vazquez *et al.*, 2016; Hanahan and Weinberg, 2011). Such changes highlight gene dysregulation in the context of explaining heterogeneous mechanisms and treatment avenues (Taylor *et al.*, 2009).

Characterizing the molecular underpinnings that associate with survival of patients with various cancers (Schramm, Li, *et al.*, 2013) as opposed to only distinguishing the differences between tumour and normal tissue offers the opportunity to study the disease at a finer resolution. Importantly, this can also lead to an increased accuracy of prediction of patient prognosis (Barter *et al.*, 2014). An increased understanding of the differences in gene coordination among patients across an important biological variable like survival outcome will improve understanding of disease mechanism and potential treatment options.

Methods exist for determining differences in correlation between two distinct groups of patients (Schramm, Li, *et al.*, 2013; Siska and Kechris, 2017; Siska *et al.*, 2016; Lai *et al.*, 2004; Fukushima, 2013), the most relevant of which is Fisher’s Z-transformation to test for differential correlation, described further in (Fukushima, 2013). However, these rely on first distinguishing two distinct sample groups for comparison. In the case where we do not have a clear way of dichotomising patients or samples, we would nevertheless like to assess some measure of differential correlation. One naïve method of determining this is to simply use a statistical linear model to test for an interaction effect between two genes with the response being the outcome of interest (such as survival time or ranking of survival), indeed one might also use a survival-specific statistical model such as Cox Proportional Hazards and test for a significant interaction term between two genes. This method, along with Fisher’s Z-transformation however relies on the assumption of normally distributed gene expression, which may not be appropriate, as well as requiring samples to be dichotomised into two groups.

We have developed an approach called Differential Correlation across Ranked Samples (DCARS), which uses local weighted correlations to build a powerful and robust statistical test to identify significant variation in levels of concordance across a ranking of samples. We applied this method to 13 TCGA gene expression datasets, using all complete survival information as the basis for ranking samples. DCARS is able to identify patterns of gene coexpression across patients ordered on survival that are more complex than simple presence or absence of correlations, as well as nonlinear interactions between gene pairs. Furthermore, DCARS can be used to identify significant edges within an existing network, which can be used to interrogate, characterise and compare subnetworks across multiple cancers.

## MATERIAL AND METHODS

### DCARS (Differential Correlation across Ranked Samples) robust testing framework

The DCARS testing framework is summarised in Figure 1. For a set of samples with index i=*1,2,…I*, assume we have ranked gene expression measurements for two genes *x* and *y*, denoted by *x_i_* and *y_i_* respectively. Then for *I* sets of sample weights (Figure S1) of length *I* with (i,j)^th^ entry in W_ij_, we calculate the sequence of local weighted correlations between x and y. Weighted correlation R_xyj_ is calculated using the formula for genes *x* and *y* with a vector of weights W_ij_ for *i=1,2,…I*,

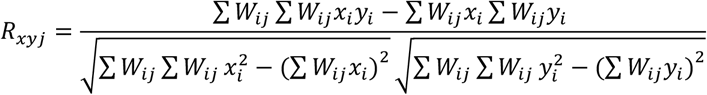

with all summations over the index *i*. The vector *R_xy_* is a sequence of weighted correlations with the weights changing akin to a sliding window, and the sample standard deviation of this sequence taken as the DCARS test statistic:

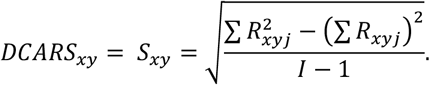

**Figure 1.**
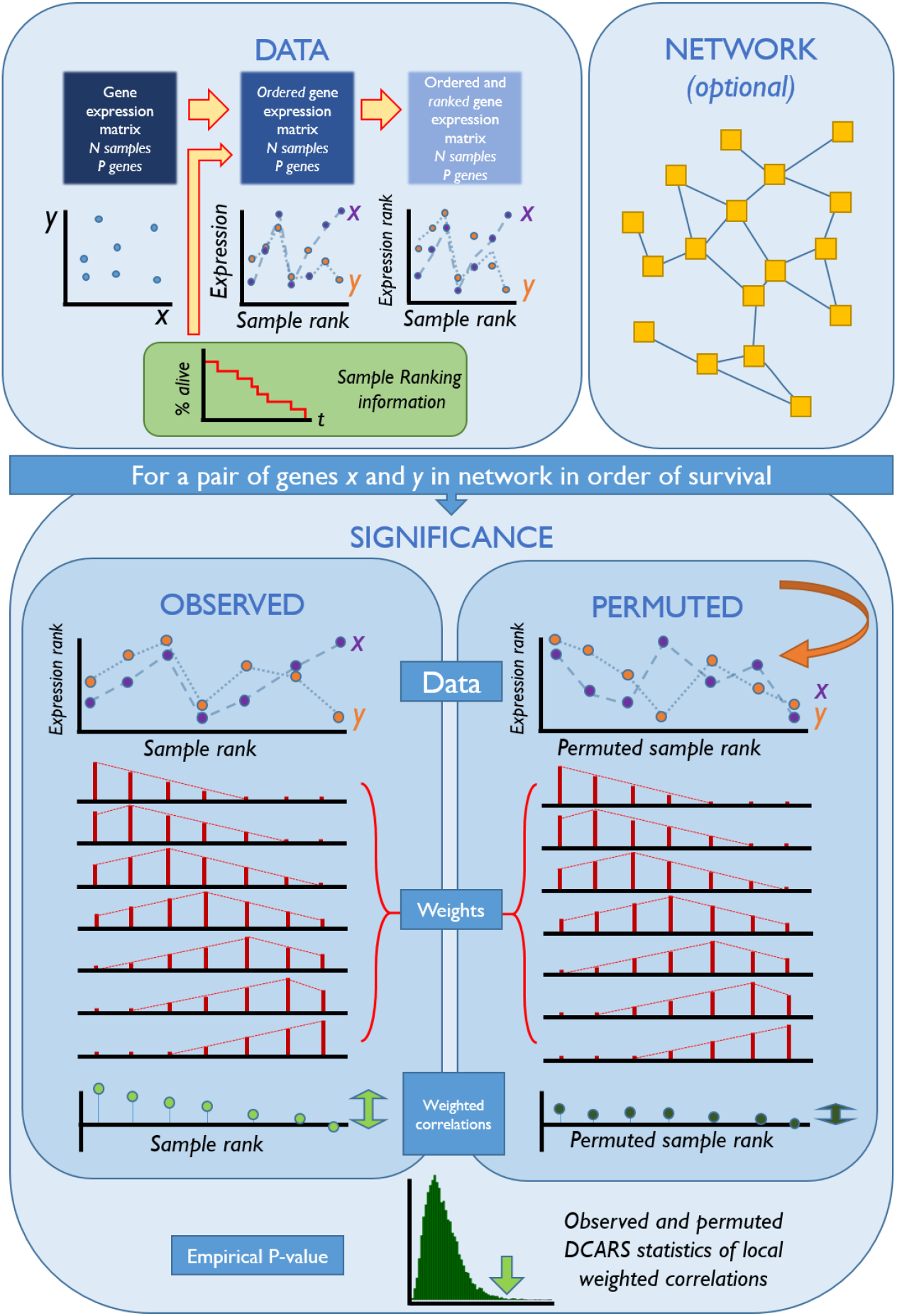
Methods flowchart. **Data**: A matrix of gene expression with N samples (columns) and p genes (rows) with accompanying complete sample ranking data, such as survival time, for the N samples is required, in which columns are ordered by survival time and gene expression values are converted to ranks for each gene across samples. **Network**: An a priori network is used to identify pairs of genes in which DCARS testing should be performed. Alternatively all paired combinations of genes can be assessed. **Significance**: To obtain the observed test statistic for a given pair of genes x and y in the network, the expression rank is taken along the survival rank. A vector of weighted correlations is calculated with a set of sample weights (triangular weights with span of 0.5 shown here), and finally the DCARS statistic is calculated as the sample standard deviation of the weighted correlation vector. Permuted DCARS statistics are obtained by rearranging the order of survival rank, ensuring that pairs of *(x,y)* expression and thus the overall correlation structure is maintained. Statistical significance is obtained by comparing the observed DCARS statistic against permuted DCARS statistics, with higher values giving evidence towards DCARS occurring for this pair of genes.

Since the null distribution of this statistics is not known at present, and may depend on factors such as the total sample size, presence of ties in the data, and choice of weight matrix, statistical significance of *S_xy_* is calculated using a permutation approach, where gene expression sample labels are repeatedly permuted at random, with higher observed values of *S_xy_* providing evidence for a difference in correlation across the sample ranking.

This method may also be used on gene expression data using the measurements themselves as opposed to the gene ranks, but to ensure robustness of the weighted correlation estimates and stability of the DCARS test statistic we ranked the gene expression measurements across samples, computing a set of quantities akin to local weighted Spearman correlation coefficients.

### Critical Values and Weight Matrices

The weighting scheme for which the local weighted correlation is calculated may vary according to different shape and parameter schemes (Figure S1), such as a triangular weight with a varying span, or a blocking weight and harmonic weight. This selection of weighting scheme affects the null distribution of the DCARS test statistic as neighbouring ranked samples are given more or less influence.

Calculating the significance of the DCARS test statistic is achieved through permutation. To reduce the running time of the testing framework across multiple pairs we first calculate the observed test statistics across all pairs of genes tested, resulting in a natural ranking of gene pairs in terms of evidence towards differential correlation along the sample ranking. The observed test statistics can be ordered to obtain a ranking or utilised as a score to assign to edges in a weighted network. To obtain statistical significance we take a stratified sample of observed test statistics and perform a permutation test on the corresponding gene pairs. This approach is used to estimate the critical value of the test statistic to obtain (unadjusted) statistical significance at a chosen level. For each of the 13 cancers listed in Table 1, we performed DCARS testing using triangular weight matrices of span 0.5, across each edge of the STRING network where both nodes existed in the cancer gene expression dataset. Further to this, we implemented a three-stage procedure which expedites the process of calculating significance as follows:

1. calculate observed DCARS test statistics,
2. calculate the 1000 permutation based *p*-values for a sample of up to 500 gene pairs stratified by the observed test statistic, and
3. loess smoothing to fit a nonlinear curve through this set of points,

thereby obtaining dataset specific critical values of the test statistic associated with a level of significance of 0.05.

**Table 1.** Details of TCGA cancer datasets. Thirteen datasets were retained for weighted correlation analysis, which had 100 or more samples with complete survival data.

| Tissue or organ type | TCGA abbreviation | Full name | Number of samples | Significant gene pairs | Number mutated samples | Number significant mutated genes |
| --- | --- | --- | --- | --- | --- | --- |
| Breast | BRCA | Breast invasive carcinoma | 151 | 725 | 134 | 60 |
| CNS | GBM | Glioblastoma multiforme | 129 | 546 | 124 | 3 |
|  | LGG | Brain Lower Grade Glioma | 125 | 728 | 123 | 10 |
| Gastrointestinal | COAD | Colon adenocarcinoma | 102 | 620 | 95 | 136 |
|  | LIHC | Liver hepatocellular carcinoma | 130 | 814 | 121 | 16 |
|  | STAD | Stomach adenocarcinoma | 145 | 481 | 144 | 55 |
| Gynecologic | OV | Ovarian serous cystadenocarcinoma | 229 | 589 | 159 | 15 |
| Head and Neck | HNSC | Head and Neck squamous cell carcinoma | 217 | 423 | 214 | 56 |
| Skin | SKCM | Skin Cutaneous Melanoma | 220 | 564 | 217 | 77 |
| Thoracic | LUAD | Lung adenocarcinoma | 183 | 1172 | 179 | 50 |
|  | LUSC | Lung squamous cell carcinoma | 212 | 375 | 207 | 74 |
| Urologic | BLCA | Bladder Urothelial Carcinoma | 178 | 706 | 177 | 49 |
|  | KIRC | Kidney renal clear cell carcinoma | 173 | 365 | 77 | 6 |

### Simulation setup

To assess the statistical properties of DCARS, we performed a simulation study. Briefly, we simulated a single pair of gene expression values *x* and *y* across 100 samples ordered by survival such that the first *j* (= 10, 30, 50, 70, 90) samples were positively correlated with a correlation coefficient of *r* (= 0, 0.25, 0.5, 0.75, 1), where *x* and *y* were simulated from bivariate normal distributions with mean 0 and variance 1. This was repeated 100 times leading to a total of 2500 simulated datasets. For each simulated dataset, values were transformed into gene ranks and each of the nine methods listed below were used to assess presence of differential correlation or association between *x* and *y*. These nine methods included

1. DCARS with triangular weight matrix with span 0.1;
2. DCARS with triangular weight matrix with span 0.5;
3. DCARS with triangular weight matrix with span 0.7;
4. DCARS with block weight matrix with span 0.1;
5. DCARS with harmonic weight matrix;
6. Linear model on original raw data testing for interaction;
7. Linear model on ranked data testing for interaction;
8. Fisher’s Z-transformation test on original raw data; and
9. Fisher’s Z-transformation test on ranked data.

Methods 1-5 were the DCARS methods, with weight matrices triangular with span 0.1, 0.5, and 0.7, block with span 0.1, and harmonic (Figure S1 shows these weights); method 6-7 are two linear model methods where the sample ranking was used as the response to a multiple linear regression with interaction between *x* and *y* with *p*-values associated with the interaction term considered the result, for the original simulated data and for the ranked data. Methods 8-9 were Fisher’s Z-transformation test for differential correlation, where the 100 samples were split into the first 50 and the last 50, utilising the raw and ranked data respectively. For the DCARS methods *p*-values were calculated using 100 permutations, and the performance of the statistical test was assessed by calculating the proportion of repetitions in which the test was called significant (*p*-value < 0.05). Based on the results of this simulation, we were able to determine the statistical properties of DCARS as well as identify suitable values for the corresponding weight matrices.

### TCGA data processing

Gene expression (FPKM), somatic mutation, and associated clinical data from The Cancer Genome Atlas (TCGA) were downloaded using the TCGAbiolinks R package (Colaprico *et al.*, 2016), specifically the Data release 8.0 of August 22 2017. Complete survival data was used, i.e., censored values were removed for this analysis. We then chose to continue with cancer datasets that had at least 100 complete survival data points, resulting in a mean of 168 samples across 13 datasets, across eight tissue or organ types including breast, central nervous system, gastrointestinal, gynecologic, head and neck, skin, thoracic and urologic tissues, summarised in Table 1. Gene expression data was filtered so that there was at least 1 FPKM for at least 20% of observations, for which a mean of 15,480 genes passed filtering. Furthermore, genes were intersected with the nodes in the STRING protein-protein interaction database (Szklarczyk *et al.*, 2015) resulting in a mean of 4484 genes across the 13 cancer datasets. Somatic mutation information was curated across genes where a sample and gene was scored as 1 if a non-silent mutation existed in that gene for that sample. This set of information was overlapped with the STRING database resulting in a mean of 3,327 mutated genes being queried across the 13 TCGA datasets.

### A priori information curation

#### STRING protein-protein interaction network

Utilising a priori networks is useful for identifying patterns of gene expression that are more likely to be causative, as well as reducing computational burden (Schramm, Jayaswal, *et al.*, 2013; Ghazanfar and Yang, 2016). Thus in order to meaningfully assess gene pairs, we used the STRING protein-protein interaction database (Szklarczyk *et al.*, 2015). The raw STRING-DB information (version 10.5) was downloaded and restricted to human (code 9606) interactions. High confidence interactions were chosen to be interactions with a score of 980 or above. This resulted in a network containing 23,678 edges across 5,508 nodes.

#### Known Cancer Gene Lists

In order to evaluate the associations of identified genes with cancer we downloaded the list of genes from the Cancer Gene Census (CGC) (Futreal *et al.*, 2004) on Jan 19 2018, containing a list of 699 genes implicated in cancer. In addition, we downloaded the list of genes from the Network of Cancer Genes version 5.0 (An *et al.*, 2016), a manually curated list of 1,571 protein coding cancer genes. The union of these two sets of cancer related genes was taken as the final list of ‘known’ cancer related genes.

#### Gene Pathway Information

To further assign biological meaning to the identified genes and sets of genes, we utilised the REACTOME pathways from the C2 curated gene sets collection obtained from the Molecular Signatures Database (MSigDB, version 5.1) (Liberzon *et al.*, 2015). For specific known gene sets we assigned the most representative pathways to them by first selecting the smallest pathway with the highest proportion of representation of the given gene set. This method results in any gene set being given an annotation, so long as at least one gene in the gene set belongs to any one of the pathways.

### Testing and estimation of DCARS genes using TCGA data and STRING

We tested for DCARS by calculating the test statistics across all edges of the STRING network and selecting gene pairs with unadjusted *p*-value of 0.05. To simplify interpretation, we then characterised these gene pairs in terms of the weighted correlation vector across the sample ranking, i.e. gene pairs were classed as positive, negative or zero on the low survival end if their mean weighted correlation across the first 10% of samples were between 0.3 and 1, −0.3 and 0.3, and between −1 and −0.3 respectively. Similarly gene pairs were classed as positive, negative or zero on the high survival end if their mean weighted correlation across the last 10% of samples were between 0.3 and 1, −0.3 and 0.3, and between −1 and −0.3 respectively. These sets of significant gene pairs formed networks and were interrogated further for biological insight.

### Testing for association of somatic mutations across sample rankings

In order to facilitate interpretations of results associated with the gene expression data, we also performed a Wilcoxon Ranked Sum Test with the matching somatic mutation data across survival rankings, specifically asking if the presence of at least one non-silent somatic mutation was associated with survival ranking. We performed this per cancer using the intersection of samples in which the gene expression data, complete survival and mutation information were available, selecting genes with an unadjusted *p*-value below 0.05 as having a gene whose mutation is significantly associated with survival ranking. The number of samples used in each cancer is given in Table 1.

### Evaluation of DCARS based on TCGA data

We used the Cancer Gene Census (CGC) and Network of Cancer Genes 5.0 (NCG) lists of genes to evaluate if genes had previously been associated with cancer. In particular we performed a one-sided Wilcoxon Rank Sum test to assess whether there was a positive association between gene pairs that contained at least one gene in the known cancer gene list and DCARS test statistics for that gene pair. As a comparison, we also performed Fisher’s Z-transformation test on the samples split 50/50 on the ranked gene expression data as described in the Simulation setup section.

## RESULTS

### Differential Correlation across Ranked Samples (DCARS) enables efficient identification of significant gene pairs

Our novel DCARS approach identifies gene pairs that show differential correlation across patient survivals. We applied our approach to each of the 13 cancers listed in Table 1 across each edge of the STRING network where both nodes existed in the cancer gene expression dataset (Supplementary File 1). Using the DCARS method, we identified a total of 8,108 unique significant (unadjusted p-value 0.05) gene pairs across the 13 TCGA datasets, which were further characterised as having changes in correlation between positive, zero and negative across the survival rankings (Figure S2). We found that the number of significant gene pairs was between 365 (KIRC) and 1,172 (LUAD) with a median of 589 (Table 1). Among these 8,108 gene pairs we found that there was a set of 1,305 unique gene pairs that were shared among at least two cancers, corresponding to 1,349 genes.

### DCARS gene pairs highly enriched with known cancer genes when applied to TCGA data

Compared to the typical Fisher Z-transformation test for differential correlation and the linear model with test for interaction term, DCARS ranked gene pairs containing known cancer genes more highly when applied to TCGA data. A significant association between the known cancer gene list and DCARS gene pair ranking was observed for 7 of the 13 cancers studied (Figure 2A), whereas only 1 and 3 cancers were associated with known cancer genes using the Fisher Z-transformation test and linear model interaction test respectively. This is despite having observed a moderate positive association between Fisher’s Z-transformation test −log10(*p*-values) and DCARS test statistics across all cancers, with Spearman correlations between 0.39 and 0.59 (Figure S3), and similarly for the linear model interaction test (between 0.47 and 0.72 Figure S4).

**Figure 2.**
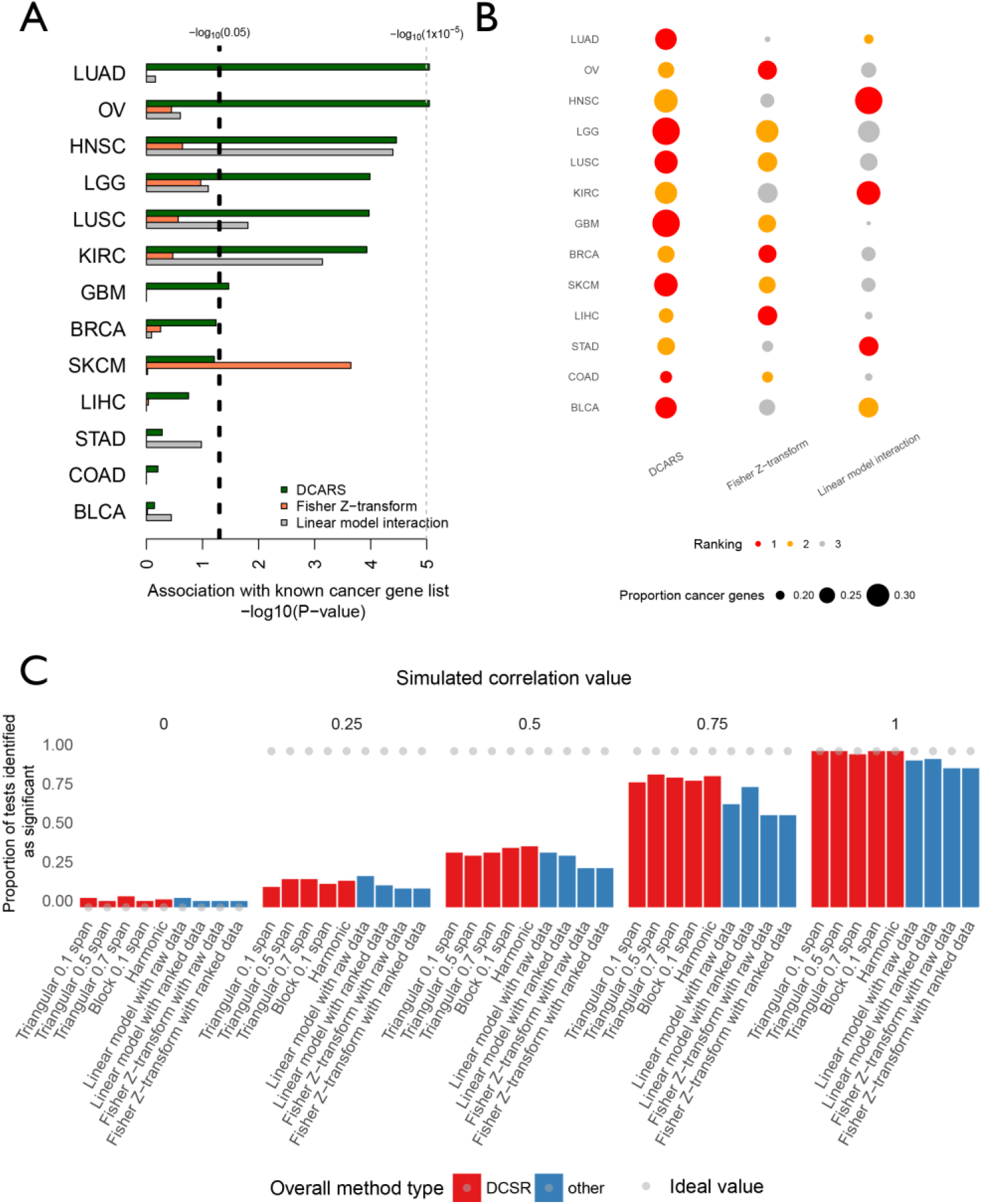
Comparison of DCARS and other statistical tests with TCGA data and in simulation. **A.** Barplot of associations with the known cancer gene list using DCARS, Fisher’s Z-transformation test, and the linear model interaction test. We performed a one-sided Wilcoxon Rank Sum test to assess if a positive association between gene pairs containing at least one gene in the known cancer gene list existed with the resulting gene pair ranking determined by DCARS (test statistic), Fisher Z-transformation test −log(*p*-value), and linear model interaction test −log(*p*-value) respectively. Barplots show the −log(*p*-value) of the Wilcoxon Rank Sum test, with vertical line corresponding to a *p*-value of 0.05, bars are capped at *p*-value of 1×10^−5^. Seven of the 13 TCGA datasets are significant using DCARS, whereas only three are significant using Fisher’s Z-transformation test and a single cancer significant for the linear model interaction test. **B.** Dotplot of proportion of known cancer genes represented in the top 1000 gene pairs for each cancer and each method. Dots are coloured by their ranking with red dots corresponding to the method for which the highest proportion of cancer genes appear in the top gene pairs list, followed by orange and then grey dots. The size of dots corresponds to the proportion of known cancer genes appearing in the top gene pairs list. DCARS is ranked first for seven cancer datasets and ranked second in the remaining six datasets. **C.** Barplots show the proportion of tests for simulated data that were considered significant with a *p*-value of 0.05 across a set of 100 data points where the first set of 30 values were correlated at values of 0, 0.25, 0.5, 0.75 and 1. Grey dots show the ideal values associated with a false positive rate of 0 and statistical power of 1. Simulations with correlated values of 0 adhered to the null model of no differential correlation across data points and thus estimate the false positive rate. DCARS methods perform similarly to other statistical tests in terms of estimated false positive rate. DCARS methods either perform similarly or outperform other statistical tests in other simulation scenarios, e.g. when the number of first values correlated is far from 50/100, and when simulated correlation values are 0.5 or less.

An additional measure of propensity of recapturing known cancer genes is to identify the proportion of known cancer genes that appear in the top ranked list of gene pairs, using the DCARS, Fisher Z-transformation test and the linear model interaction test. We calculated the proportion of known cancer genes appearing in the top 1,000 gene pairs for each TCGA dataset and each method, and found that DCARS consistently selected a higher proportion of known cancer genes than the other two methods, with the highest proportion for seven cancer datasets and the second highest for the remaining six datasets (Figure 2B, Figure S6).

Having identified a strong overrepresentation of gene pairs with DCARS and at least one gene belonging to the known cancer gene list provides evidence that the identified gene pairs are strongly of biological interest and DCARS is a useful method at identifying gene relationships of interest, as well as highlighting potential novelty of suggested mechanisms of known cancer genes.

### DCARS statistics are robust to choice of weights leading to computational efficiency

Monotonic relationships were found between observed test statistics and the corresponding permutation based *p*-values (Figure S6). Furthermore, critical values were found to be dependent on the number of samples in the dataset and the choice of weighting scheme (Figure S7A). The DCARS test statistic is influenced by the choice of weights, however if the same weighting scheme is used, then the distribution of these test statistics is similar across cancers and depends on the sample size (Figure S7A). We found that the observed critical values for the TCGA datasets varied between 0.098 (OV) and 0.15 (COAD) at a significance level of 0.05. Further interrogation revealed there was an overall monotonic trend between the number of samples in the dataset and the estimated DCARS critical value (Figure S7B). Randomly subsetting from the largest dataset (OV with 229 samples) and repeating the analysis confirmed that the critical value was indeed related to the sample size (Figure S8 and Figure S9). The estimated critical value is also dependent on the choice of weight matrix used. Repeating the OV subset analysis with a triangular weight matrix of different span of 0.1 (Figure S9), revealed that the curve between sample size and the corresponding critical values is roughly parallel to that when using a triangular weight matrix of span 0.5 (Figure S7C). Importantly, we also found that the ranking of test statistics was mostly preserved using either a triangular weight matrix of span 0.1 or 0.5 (Figure S7D), suggesting DCARS is fairly robust to somewhat arbitrary choice of weighting spans. Thus we suggest that utilising a triangular matrix of span 0.5 is a suitable choice for weight matrix.

### DCARS is a more powerful test especially with imbalanced groups of correlated samples

While DCARS identifies more cancer related associations than other typical differential correlation approaches, we also show that DCARS is more powerful through simulation (Figure 2C and Figure S10). Overall, DCARS methods either performed similarly or outperformed other statistical tests for identifying changes in correlation associations between *x* and *y*. DCARS was consistently more powerful than the linear model with interaction test. The Fisher Z-transformation test method first split samples into the first 50% and last 50% and tested for differential correlation. In the case of correct model specification (third row of Figure S10), where the first set of 50 samples were simulated with positive correlation, the Fisher Z-transformation test was marginally more powerful, but in the case of model misspecification (for example shown in Figure 2C), DCARS maintained a higher level of power. These simulation results highlight the ability of DCARS to powerfully capture differences in correlation robustly without needing to specify two sets of samples.

In terms of characterising false positive error, simulating a correlation value of zero corresponds to the null model, where in fact no difference in correlation exists across samples (leftmost column in Figure S10). Thus, estimates of proportion of significance for this simulation setting are estimates of the false positive error rate. DCARS methods performed similarly to the other methods in terms of the estimated false positive error rate (Figure S11), with median estimated false positive error rates ranging between 0.04 and 0.07 for the DCARS methods and 0.04 and 0.06 for the other statistical tests, indicating that DCARS controls the false positive rate while maintaining statistical power.

### DCARS captures changes in correlation that are conserved across multiple cancers

In order to identify gene pairs that were associated with global patterns of gene expression in cancer, we looked specifically at gene pairs that were significant and consistent across multiple cancers. The REACTOME pathways (C2 from MSigDB) were used to further explore biological processes associated with the genes we identified. We selected gene pairs that were 1) significant and 2) had weighted correlation patterns in a consistent direction. We identified three gene pairs consistently significant across four cancers (Table 2), 19 that were significant across three cancers (Table 2) and a further 430 pairs that were significant over two cancers (Supplementary File 2). In particular at least one the three specific gene pairs, COPA and COPE, that had this pattern across four cancers (Figure 3A) has previously been shown to be associated with biological processes in cancer, discussed further below.

**Table 2.**
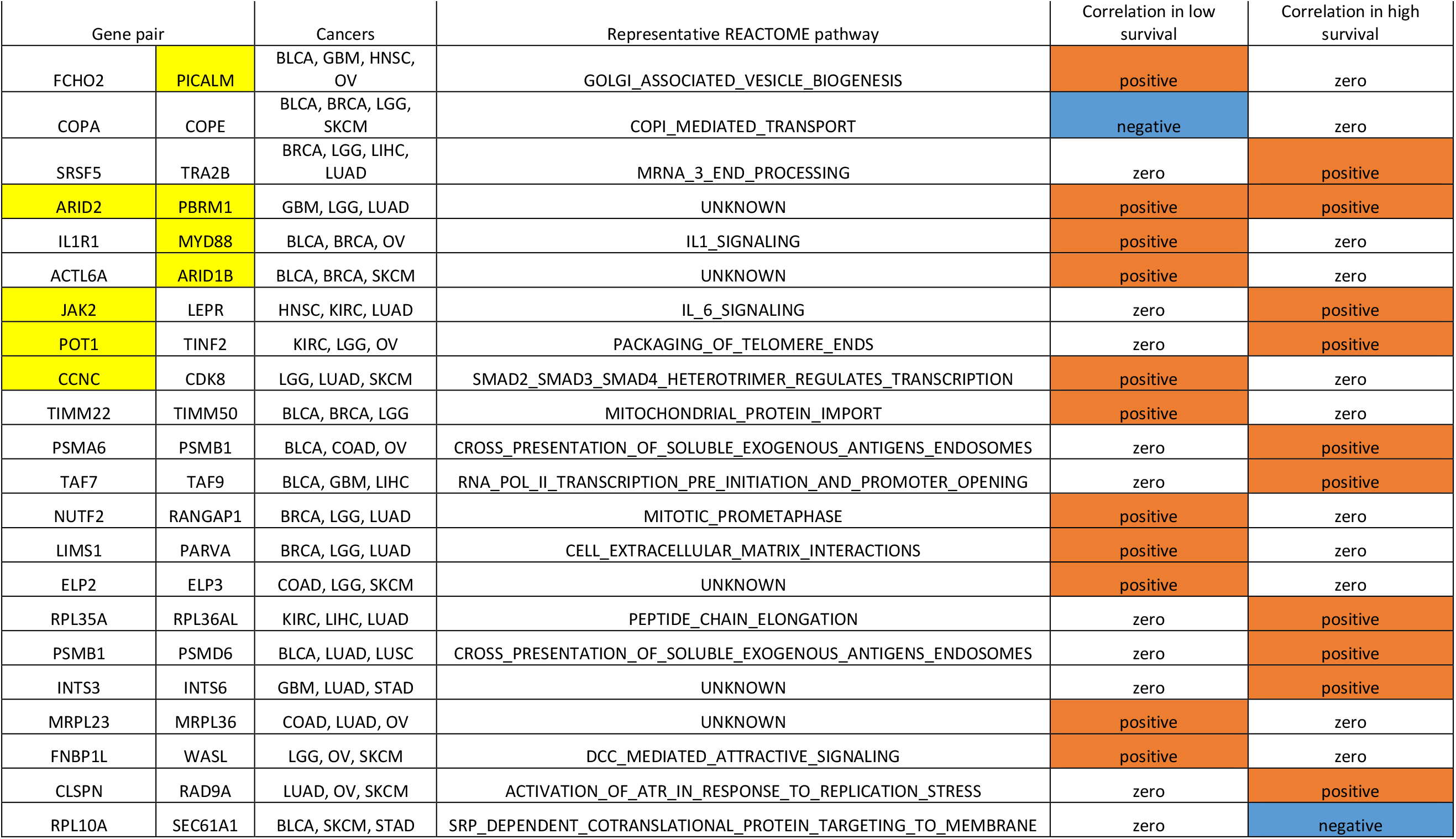
Significant gene pairs across three or more cancers. Gene pairs were filtered so that they were significant in the same direction in at least 3 cancers. Highlighted genes belong to the list of known cancer genes, gene pairs are sorted alphabetically, with most representative REACTOME pathway given. Correlation was identified as zero, negative, or positive at the low or high end of survival samples.

**Figure 3.**
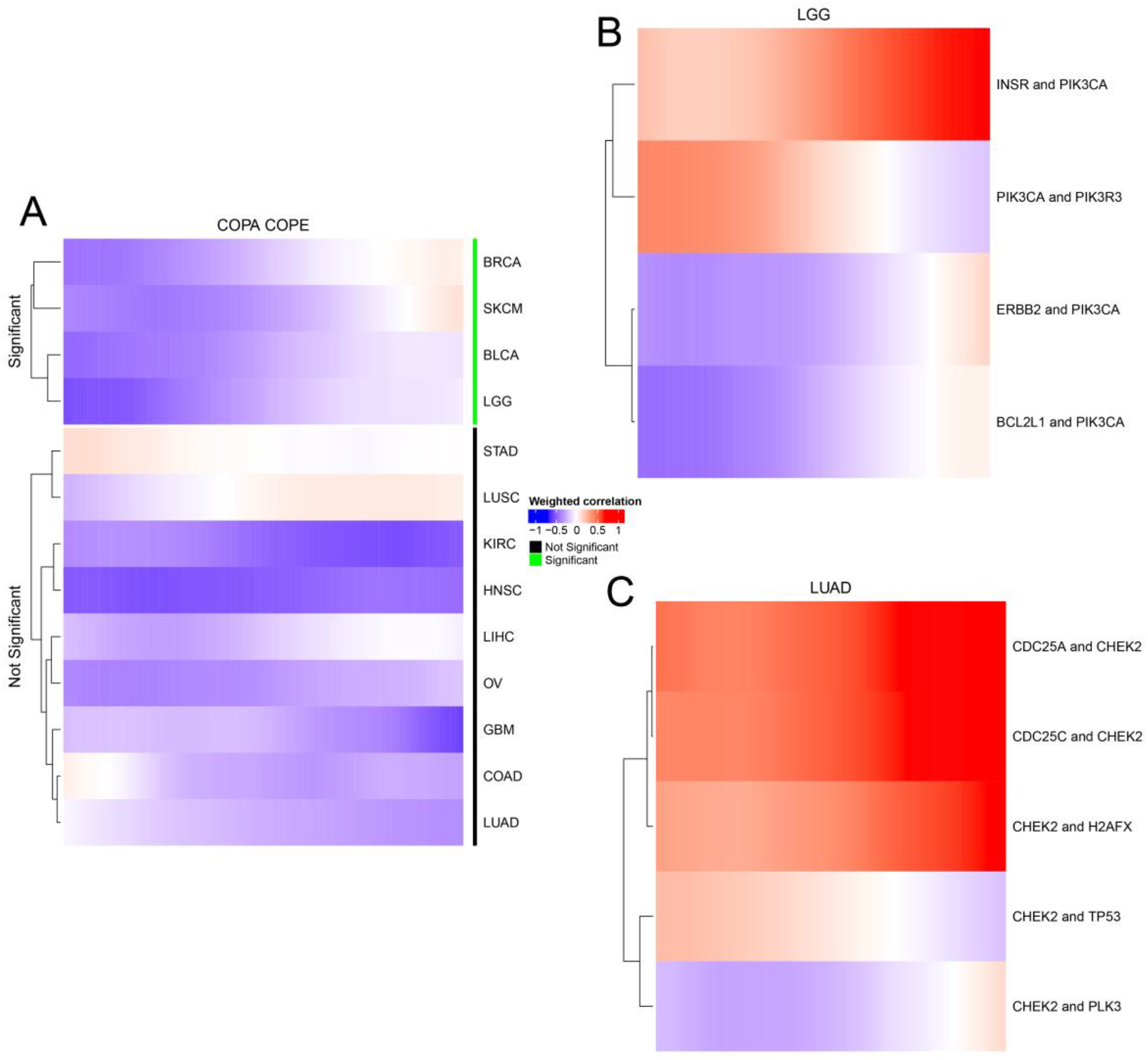
Weighted correlation vectors of specific gene pairs of interest. **A.** Local weighted correlation vectors from low (left) to high (right) survival of COPA and COPE across all 13 TCGA datasets. Green bars on the side indicate significant DCARS gene pairs. **B.** Local weighted correlation vectors from low to high survival for specific gene pairs involving PIK3CA across the LGG TCGA dataset. All rows shown are significant DCARS gene pairs. **C.** Local weighted correlation vectors from low to high survival for specific gene pairs involving CHEK2 across the LUAD TCGA dataset. All rows shown are significant DCARS gene pairs.

Of the genes belonging to gene pairs that are significant in the same direction over three or more cancers, we observed overrepresentation of genes (Supplementary File 3) belonging to processes associated with membrane budding (*p*-value<10^−4^, STRING version 10.5 functional enrichment tool (Szklarczyk *et al.*, 2015), genes include COPA, COPE, FCHO2, FNBP1L, PICALM, and WASL) and the majority of genes were involved in cellular macromolecule metabolic processes (*p*-value<10^−3^, highly connected genes include JAK2, SEC61A1, RPL35A, among other genes such as CLSPN, RAD9A, and POT1, TINF2, and PSMD6, PSMB1, and PSMA6). One identified gene pair is that of CLSPN and RAD9A found to be positively correlated in high survival patients in LUAD, OV and SKCM cancers, belonging to the TP53 pathway. RAD9 has also been implicated in cancer, where aberrant or uncontrolled expression is associated with deleterious health consequences (Lieberman *et al.*, 2011). This supports the notion that the TP53 pathway is ‘on’ when actively fighting a tumour, and suggests that tumours with low associated survival evade activation of the tumour suppressive TP53 pathway (Vogelstein *et al.*, 2000).

#### COPI-coated vesicle budding

We looked at individual gene pairs in which changes were observed in the same direction, and found a number of pairs with concordant direction of correlation (gain or loss of positive or negative correlation) across survival time. One of the most common gene pairs identified was COPA and COPE, identified to be negatively correlated in low survival and not correlated in high survival groups among BLCA, BRCA, LGG and SKCM (Figure 3A). Additionally, the LUSC network contains the gene pair COPA and COPB1 which is positively correlated in the high survival group. This set of genes is related to vesicle transport which may be recruited to carry signals across tumour cells (Wang *et al*., 2010), and thus may be associated to processes occurring in tumours.

### Network-based analysis of DCARS gene pairs allows discovery of specific pathways unique to survival groups

To demonstrate the potential of DCARS to build on meaningful biological analyses, we performed a network analysis on each of the TCGA datasets. Subnetworks of the STRING network were built by selecting the significantly DCARS gene pairs, assigning a direction of weighted correlation across survival on each edge. We were then able to interrogate the structure of the resulting networks, including identifying concordant patterns of correlation change across survival along communities detected using the walktrap algorithm implemented in the igraph R package (Csárdi and Nepusz, 2006). All figures associated with all cancers are provided in Figure S2, Figure S12, and Supplementary File 4, with those associated with a chosen cancer of skin cutaneous melanoma (SKCM) shown in Figure 4A. Rather than specifically look at the changes from positive, zero, or negative weighted correlation in the low survival group to positive, zero, or negative weighted correlation in the high survival group, we instead noted for gene pairs if there was one of 1) correlation of any direction present in the low survival group and not in the high survival group, 2) correlation of any direction present in the high survival group and not in the low survival group, or 3) if there was a switch in correlation direction between the high and low survival group. This approach revealed a number of distinct subnetworks within the DCARS derived network contained a high proportion of edges with correlations present in either the low or high surviving group (Figure 4B and Figure 4C). This suggests that DCARS can point towards processes that may be switched ‘on’ in one group of samples and ‘off’ in the other. One such example in the SKCM DCARS network is the subnetwork associated with Cell Cycle Checkpoints, where the majority of correlation is present in the high survival group (Figure 4D), corroborating with the notion that processes associated with cell cycle are altered in melanoma (Kaufmann *et al.*, 2008). This specific characterisation of subnetworks in terms of identifying correlation in the low or high survival set of samples allows for deeper understanding of the biological mechanisms at play in terms of the hallmarks of cancer (Hanahan and Weinberg, 2011), and potentially lead towards identification of therapeutic targets.

**Figure 4.**
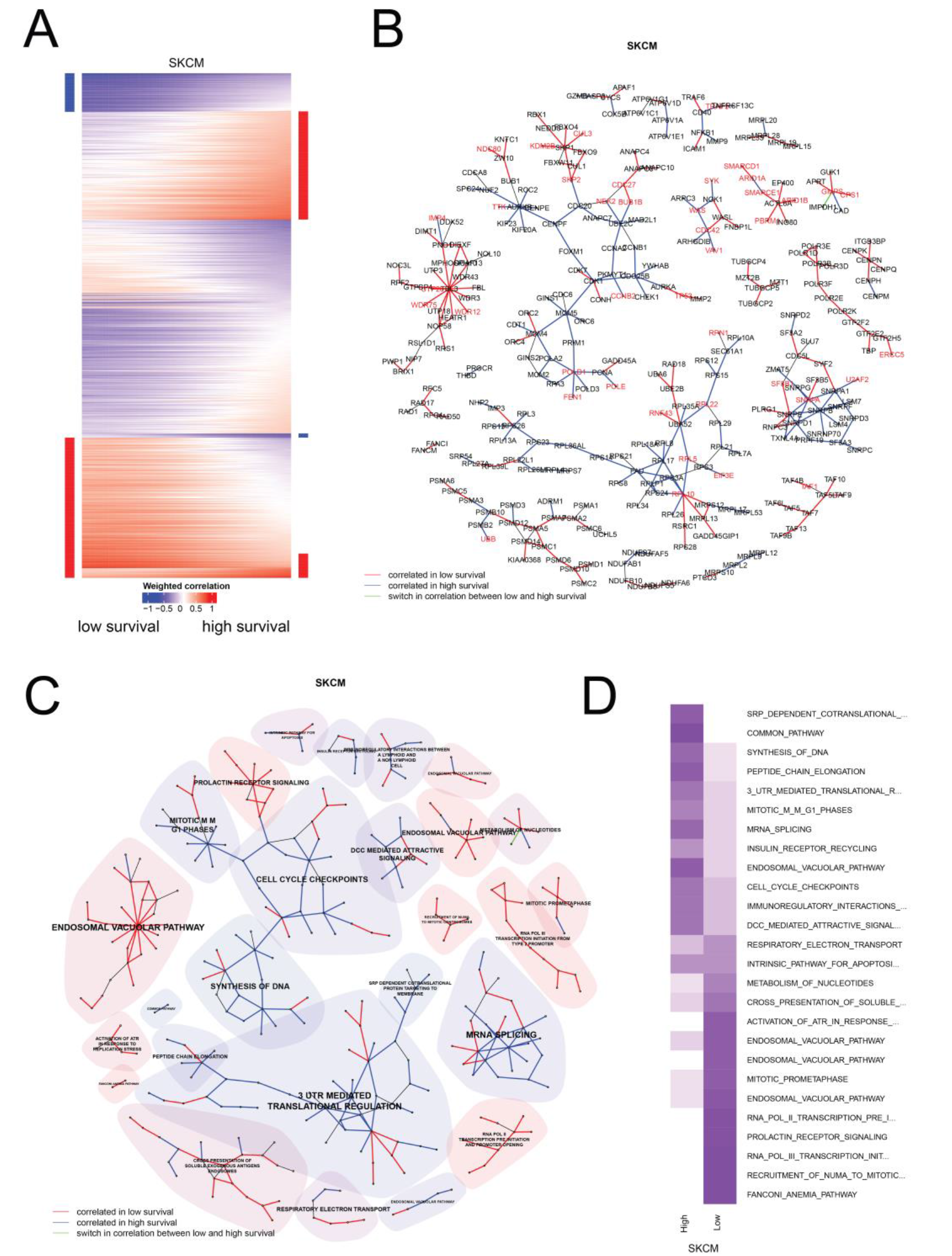
DCARS analysis results for skin cutaneous melanoma (SKCM) TCGA dataset. **A**. Heatmaps display the weighted correlation vectors from low to high survival samples, with red indicating positive weighted correlation and blue negative weighted correlation. Bars along the sides indicate gene pairs for which the weighted correlation is identified as positive (red) or negative (blue) or zero (white) at either low or high end of survival, thus characterising each gene pair in terms of the direction of change of weighted correlation. **B**. Subnetwork of STRING network generated by obtaining DCARS gene pairs, where disjoint subnetworks with fewer than 5 nodes removed for clarity, red edges represent gene pairs correlated in low survival samples, blue edges represented gene pairs correlated in high survival samples, green pairs represent a switch in correlation between low and high survival samples. **C**. Subnetwork as in panel B with community detection result overlaid, and most representative REACTOME pathway labelled. Shading indicates the proportion of edges representing gene pairs correlated in low survival (red) and in high survival (blue). **D**. Heatmap of proportion of gene pairs attributed to either high (correlated in high survival) or low (correlated in low survival) group for each community shown in panel C, labelled by the most representative REACTOME pathway.

### Integrating DCARS with somatic mutation information points to potentially causative biological associations

We utilised the TCGA somatic mutation information, by identifying genes with nonsilent mutations that associated with survival ranking (Figure S13), taking into account whether gene mutations were associated with a lower or a higher survival ranking. We then associated the gene mutation results with DCARS gene pairs by identifying significant gene pairs that contained a significantly mutated gene. We further split these gene pairs of interest based on 1) if the mutated gene is associated with high or low survival ranking, and 2) if at least one of the gene pair appears in the list of known cancer genes. These lists of mutated genes and gene pairs are given in Table 3 (contain known cancer genes and mutated in low survival samples), Table S1 (contain known cancer genes and mutated in high survival samples), Table S2 (do not contain known cancer genes and mutated in low survival samples), and Table S3 (do not contain known cancer genes and mutated in high survival samples). We identified 9 known cancer genes that were more mutated in the low surviving group and also belonged to a gene pair that was DCARS for the same cancer. In addition to these 8 genes, we also found 28 genes not in the known cancer gene list that were more mutated in the low surviving group and also belong to a gene pair that was DCARS for the same cancer. We also found a total of 45 genes that were more mutated in the high surviving group with corresponding DCARS results, of which 13 appear in the known cancer gene list.

**Table 3.** DCARS gene pairs associated with somatic mutations in the low survival group with at least one known cancer gene. Highlighted yellow cells are genes that appear in the list of known cancer genes. Correlation was identified as zero, negative, or positive at the low or high end of survival samples.

| Cancer | Mutated Gene | Gene Pair |  | Low surviving correlation | High surviving correlation |
| --- | --- | --- | --- | --- | --- |
| BLCA | ESPL1 | ESPL1 | PTTG1 | zero | positive |
|  | MAP3K4 | GADD45G | MAP3K4 | negative | zero |
|  |  | GADD45B | MAP3K4 | negative | zero |
| BRCA | LYN | LYN | STAT3 | zero | positive |
|  | MSH2 | EXO1 | MSH2 | positive | positive |
|  |  | MSH2 | POLE | zero | positive |
|  |  | FEN1 | MSH2 | zero | positive |
|  | NUP214 | NUP214 | NUP85 | zero | positive |
| COAD | WRN | TOP3A | WRN | positive | zero |
| LGG | PIK3CA | BCL2L1 | PIK3CA | negative | zero |
|  |  | PIK3CA | PIK3R3 | positive | zero |
|  |  | INSR | PIK3CA | zero | positive |
| LIHC | PRKDC | PRKDC | TP53 | zero | positive |
| LUAD | CHEK2 | CDC25C | CHEK2 | positive | positive |
|  |  | CDC25A | CHEK2 | positive | positive |
|  |  | CHEK2 | H2AFX | zero | positive |
|  | KRAS | KRAS | RAF1 | zero | positive |
|  |  | BRAF | KRAS | zero | positive |
|  | NFE2L2 | HMOX1 | NFE2L2 | positive | zero |
|  | RELA | CREBBP | RELA | zero | positive |
|  | TOP2A | RANBP2 | TOP2A | zero | positive |

#### PIK3CA in LGG

The gene PIK3CA, appearing in the list of known cancer genes, was more mutated in low survival groups of lower grade glioma (LGG) (Table 3), and the corresponding DCARS LGG subnetwork in which PIK3CA belonged related to IL2 signalling (Supplementary File 4, LGG network). Within the subnetwork, correlation was found to exist between PIK3CA and ERRB2, BCL2L1, and PIK3R3 respectively in low survival samples and correlation was found to exist between PIK3CA and INSR in high survival samples (Figure 3B). Such a pattern may suggest that rather than entire processes being absent or present across one end of the set of samples, subtle switching of gene coexpression can occur in these pathways via events such as mutation. Indeed, previous literature (Karakas *et al.*, 2006) has shown that mutations in PIK3CA are associated with overactivation of the PI3K/AKT pathway leading to an increase in cell proliferation, in a number of cancers. Additionally we found this gene was also associated with COAD and SKCM with similarly more mutations observed in lower surviving patients, but no direct DCARS associations of interest were identified.

#### CHEK2 in LUAD

The gene CHEK2, a known checkpoint kinase and tumour suppressor (McGowan, 2002) and appearing in the known cancer gene list, was more mutated in low survival groups of lung adenocarcinoma (LUAD) (Table 3), and the corresponding DCARS LUAD subnetwork in which CHEK2 belonged related to Cell Cycle Checkpoint pathway (Supplementary File 4, LUAD network), specifically involving the cell cycle division genes CDC25A, CDC25C, CHEK2, and TP53. Within the subnetwork, correlation was found to exist between CHEK2 and CDC25A, CDC25C, H2AFX respectively in high survival samples and correlation was found to exist between CHEK2 and PLK2 in low survival samples, whereas a subtle switch in correlation was observed between CHEK2 and TP53 (Figure 3C). The presence of lower absolute correlation in the low survival group among these gene pairs suggests these processes are deregulated in the low survival group and may still retain typical function in the high survival group (Eymin and Gazzeri). This further analysis highlights the ability for DCARS to select gene pairs that are biologically informative and can shed light on the underlying biology.

## DISCUSSION

We have introduced a novel approach called Differential Correlation across Ranked Samples (DCARS) and have shown that it is a powerful method for identifying patterns of correlation that varies across ranked samples. This method is more flexible than typical approaches that require samples to be split into two distinct patient groups. When applied to TCGA data, our approach combined with further network analytics leads to biological insight as highlighted here with our study using TCGA data.

While we have applied DCARS to the gene expression data in TCGA, it can easily be applied to many other biological data platforms and diseases, e.g. DNA methylation patterns. DCARS is a flexible statistical approach that assesses changes in correlation of two variables across a ranking of samples. This approach is able to discover biologically informative relationships among genes across a variable of interest. Here, we have used it in the context of gene expression and survival rankings of samples, but the same approach can easily be extended to consider other biomedical data, for example identifying changes in correlation pattern between genes across single cells ordered along a pseudotime trajectory; or identifying changes in correlation between other types of clinical variables ordered along another clinical variable such as BMI. DCARS can be extended to other types of data such as categorical data, where the measure of concordance can be tailored to suit the data type. Since DCARS utilises ranked samples and transforms the gene expression data into gene ranks, it does not rely on distributional assumptions such as normality to ensure meaningful results, thus is robust to departures from normally distributed data.

Our current DCARS method requires an ordered set of samples. In the context of survival, a number of samples may be censored, and thus it is unclear what the latent or ‘true’ order of samples is. One approach to address this issue is to modify the associated weight matrix to reflect the censored sample, or to perform a repeated analysis where different plausible sample orders are used. Similar to this potential issue is the case where sample ties may occur, in which case the weight matrix may simply have equal values assigned to the tied samples.

Interestingly, there is the possibility that other factors relevant to the sample ranking can be associated with the correlation patterns observed in the data. For example, in our illustration of DCARS using cancer survival ranking, cancer samples may differ in terms of the tumour sample itself in a way that associates with survival. This could be through various factors such as the proportion of normal or stromal to tumour cells in the sample, and the level of infiltration of other cell types such as immune cells. A strategy to account for these other factors is to perform correction through a regression analysis such as those used in experimental batch correction methods, and perform DCARS on the resulting matrix of residual measurements. Another exciting possibility is to perform a cell type deconvolution algorithm to identify putative cell type compositions for the sample at hand, and then perform DCARS on the resulting cell-type specific gene expression signal across the sample ranking. This would identify changes in correlation across a sample ranking specific to a cell type of interest.

We have demonstrated the utility of the DCARS method by applying it to the TCGA gene expression dataset, whereby known cancer genes were greatly overrepresented and some interesting biological associations between cancer genes was identified. In addition we demonstrated that DCARS is an informative aspect of a network-based analysis, further enhancing existing network analytics such as community detection (Petrochilos *et al.*, 2013; Fuller *et al.*, 2007; Ideker *et al.*, 2002). In the current work we utilised a community detection approach to identify biological processes most associated with ‘presence’ or ‘absence’ of correlation on the low- or high- end of survival outcome. If coupled with corresponding gene expression of normal tissue, the ‘normal’ level of concordance between genes can be identified and thus a more informative observation be found where a pair of genes may have ‘gained’ or ‘lost’ correlation in low or high survival groups, directly associating to potential therapeutic targets.

In summary, our novel Differential Correlation across Ranked Samples (DCARS) method is a powerful and flexible testing framework for identifying changes in correlation patterns across a ranking of samples, allowing for deep interrogation of the complex relationships observed in gene expression systems.

## AVAILABILITY

Publicly available data from The Cancer Genome Atlas (TCGA) was used using the TCGABiolinks R package (Colaprico *et al.*, 2016). Supplementary Files and software for running DCARS using R is available at https://github.com/shazanfar/DCARS.

## AUTHORS’ CONTRIBUTIONS

SG conceived the study with input from EP, JYHY and JTO. SG performed the data analysis in R and wrote the manuscript with input from EP on the design, analytics, interpretation and the direction of the study. DS curated and processed all the TCGA datasets. All authors read and approved the final version of the manuscript.

## ACKNOWLEDGEMENT

The authors thank all their colleagues, particularly at The University of Sydney, School of Mathematics and Statistics, for their support and intellectual engagement. The following sources of funding for each author, and for the manuscript preparation, are gratefully acknowledged: Australian Research Council Discovery Project grant DP170100654 (JYHY, JTO, DS); Australia. NHMRC Career Developmental Fellowship APP1111338 (JYHY); School of Mathematics and Statistics, The University of Sydney (EP, SG); and the Judith and David Coffey Life Lab at the Charles Perkins Centre, The University of Sydney (SG). The funding source had no role in the study design; in the collection, analysis, and interpretation of data, in the writing of the manuscript, and in the decision to submit the manuscript for publication.

---

**Table S1.** DCARS gene pairs associated with somatic mutations in the high survival group with at least one known cancer gene. Highlighted cells are genes that appear in the list of known cancer genes. Correlation was identified as zero, negative, or positive at the low or high end of survival samples.

| Cancer | Mutated Gene | Gene Pair |  | Low surviving correlation | High surviving correlation |
| --- | --- | --- | --- | --- | --- |
| BLCA | MYH9 | MYH9 | MYL9 | zero | positive |
|  | PTK2 | PIK3CA | PTK2 | positive | zero |
| COAD | DCTN1 | DCTN1 | DCTN2 | negative | zero |
|  | NUP88 | NUP214 | NUP88 | negative | zero |
| HNSC | TSC2 | TSC1 | TSC2 | zero | positive |
| LUSC | CDC27 | ANAPC13 | CDC27 | positive | zero |
|  | TP53 | SUMO1 | TP53 | zero | positive |
|  |  | SP1 | TP53 | zero | positive |
|  |  | SIRT1 | TP53 | zero | positive |
| OV | KAT6B | BRPF1 | KAT6B | positive | zero |
|  |  | BRD1 | KAT6B | positive | zero |
|  | MLH1 | MLH1 | PCNA | zero | positive |
| SKCM | BRCA1 | ATM | BRCA1 | positive | zero |
|  | RPTOR | RPTOR | RRAGB | negative | zero |
|  |  | PRKAA1 | RPTOR | negative | zero |
|  | RSRC1 | RPL10 | RSRC1 | negative | zero |
|  | XPO1 | RANGAP1 | XPO1 | negative | zero |
| STAD | IRS1 | INSR | IRS1 | zero | positive |
|  | MUC1 | MUC1 | MUC2 | zero | positive |
|  | MYD88 | MYD88 | TLR8 | positive | zero |
|  |  | MYD88 | TLR4 | positive | zero |
|  |  | MYD88 | TLR1 | positive | zero |
|  |  | MYD88 | TLR6 | positive | zero |

**Table S2.**
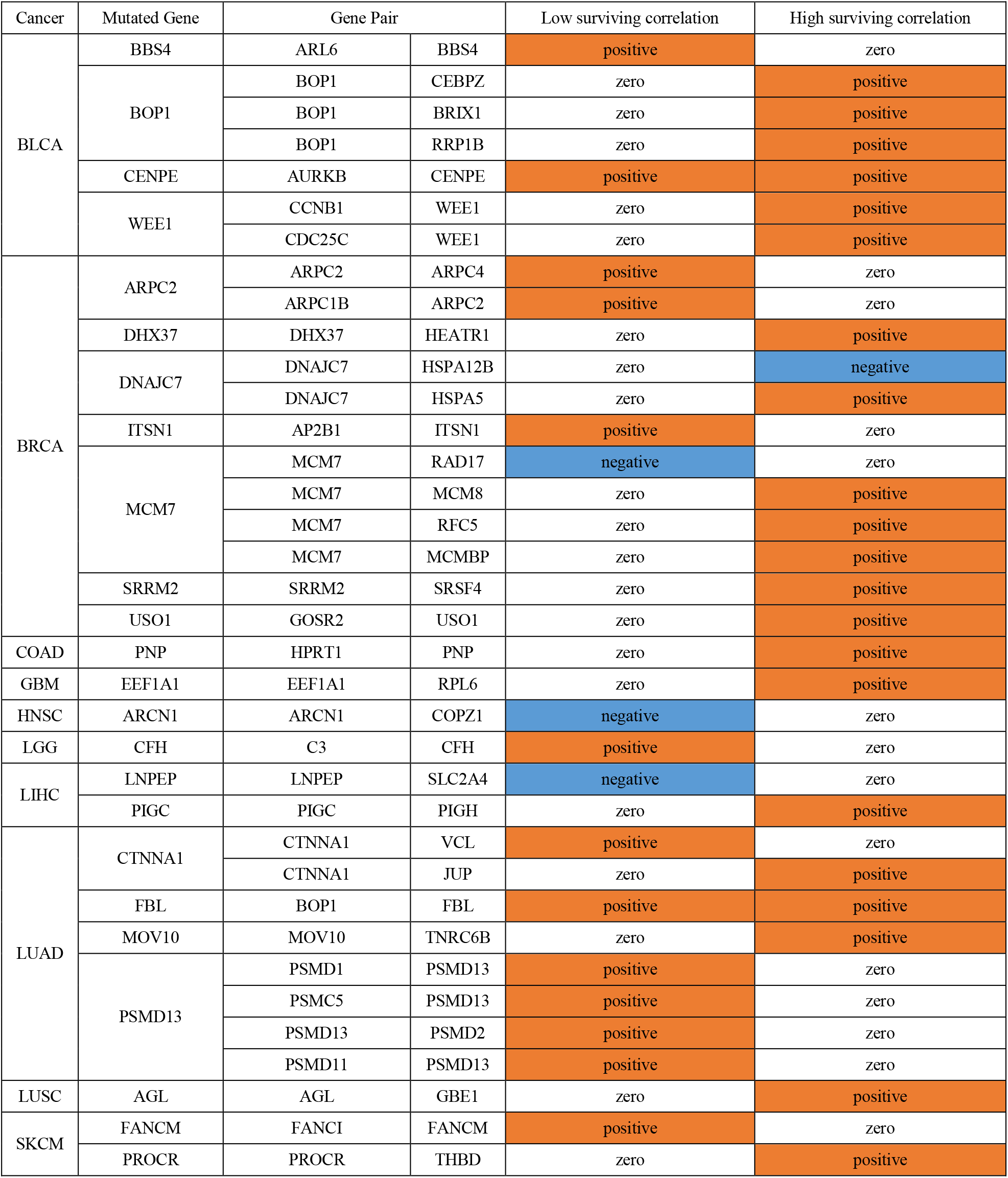
DCARS gene pairs associated with somatic mutations in the low survival group with no known cancer genes. Correlation was identified as zero, negative, or positive at the low or high end of survival samples.

**Table S3.** DCARS gene pairs associated with somatic mutations in the high survival group with no known cancer genes. Correlation was identified as zero, negative, or positive at the low or high end of survival samples.

| Cancer | Mutated Gene | Gene Pair |  | Low surviving correlation | High surviving correlation |
| --- | --- | --- | --- | --- | --- |
| BLCA | PPP1R12A | PPP1CC | PPP1R12A | positive | zero |
|  | UCLH5 | PSMB10 | UCLH5 | negative | zero |
|  |  | PSMD8 | UCLH5 | zero | positive |
|  |  | PSMB7 | UCLH5 | zero | positive |
|  |  | PSMB4 | UCLH5 | zero | positive |
|  |  | PSMB2 | UCLH5 | zero | positive |
|  |  | PSMA3 | UCLH5 | zero | positive |
|  |  | PSMA4 | UCLH5 | zero | positive |
|  |  | PSMD3 | UCLH5 | zero | positive |
|  |  | PSMA2 | UCLH5 | zero | positive |
| COAD | COPG1 | COPB1 | COPG1 | zero | negative |
|  | DHX37 | DHX37 | WDR36 | zero | positive |
|  | DLD | DLD | GCSH | zero | positive |
|  | EXOC3 | EXOC3 | EXOC7 | zero | positive |
|  | EXOSC4 | EXOSC1 | EXOSC4 | positive | zero |
|  |  | EXOSC2 | EXOSC4 | positive | zero |
|  | FZD7 | FZD7 | WNT5B | zero | positive |
|  | GEMIN5 | GEMIN5 | SNRPD1 | positive | zero |
|  | MUT | ACLY | MUT | negative | zero |
|  | NFS1 | LYRM4 | NFS1 | zero | positive |
|  | POLR2D | GTF2B | POLR2D | zero | positive |
|  |  | POLR2C | POLR2D | zero | positive |
|  | PRPF8 | PRPF8 | TXNL4A | zero | positive |
|  | RPL19 | RPL15 | RPL19 | zero | positive |
|  | RPS6KB2 | RPS6 | RPS6KB2 | positive | zero |
|  | SMARCC1 | ACTL6A | SMARCC1 | zero | positive |
|  |  | SMARCC1 | SMARCD2 | zero | positive |
|  | SUPT16H | CSNK2A1 | SUPT16H | positive | zero |
|  | TRADD | TRADD | TRAF2 | zero | positive |
| HNSC | COG4 | COG4 | COG6 | negative | zero |
|  | PRPF6 | LSM3 | PRPF6 | positive | zero |
| KIRC | BMS1 | BMS1 | RCL1 | positive | zero |
| LUAD | DIEXF | DIEXF | UTP14C | zero | positive |
|  |  | DIEXF | UTP3 | zero | positive |
|  |  | DIEXF | NOL6 | zero | positive |
|  | MRPL14 | MRPL14 | MRPL22 | zero | positive |
|  | STAM | STAM | UBC | positive | zero |
| LUSC | CKAP5 | CKAP5 | TACC2 | positive | zero |
|  | IQGAP2 | CDC42 | IQGAP2 | zero | positive |
|  | PRPF8 | DHX15 | PRPF8 | zero | positive |
|  | XAB2 | CWC22 | XAB2 | positive | zero |
| SKCM | RSRC1 | RPS28 | RSRC1 | negative | zero |
| STAD | GABARAPL1 | ATG4A | GABARAPL1 | negative | zero |
|  |  | BECN1 | GABARAPL1 | negative | zero |
|  | MRPL16 | GFM1 | MRPL16 | positive | zero |
|  |  | MRPL16 | MRPL34 | zero | positive |
|  |  | MRPL16 | MRPL43 | zero | positive |

**Figure S1.**
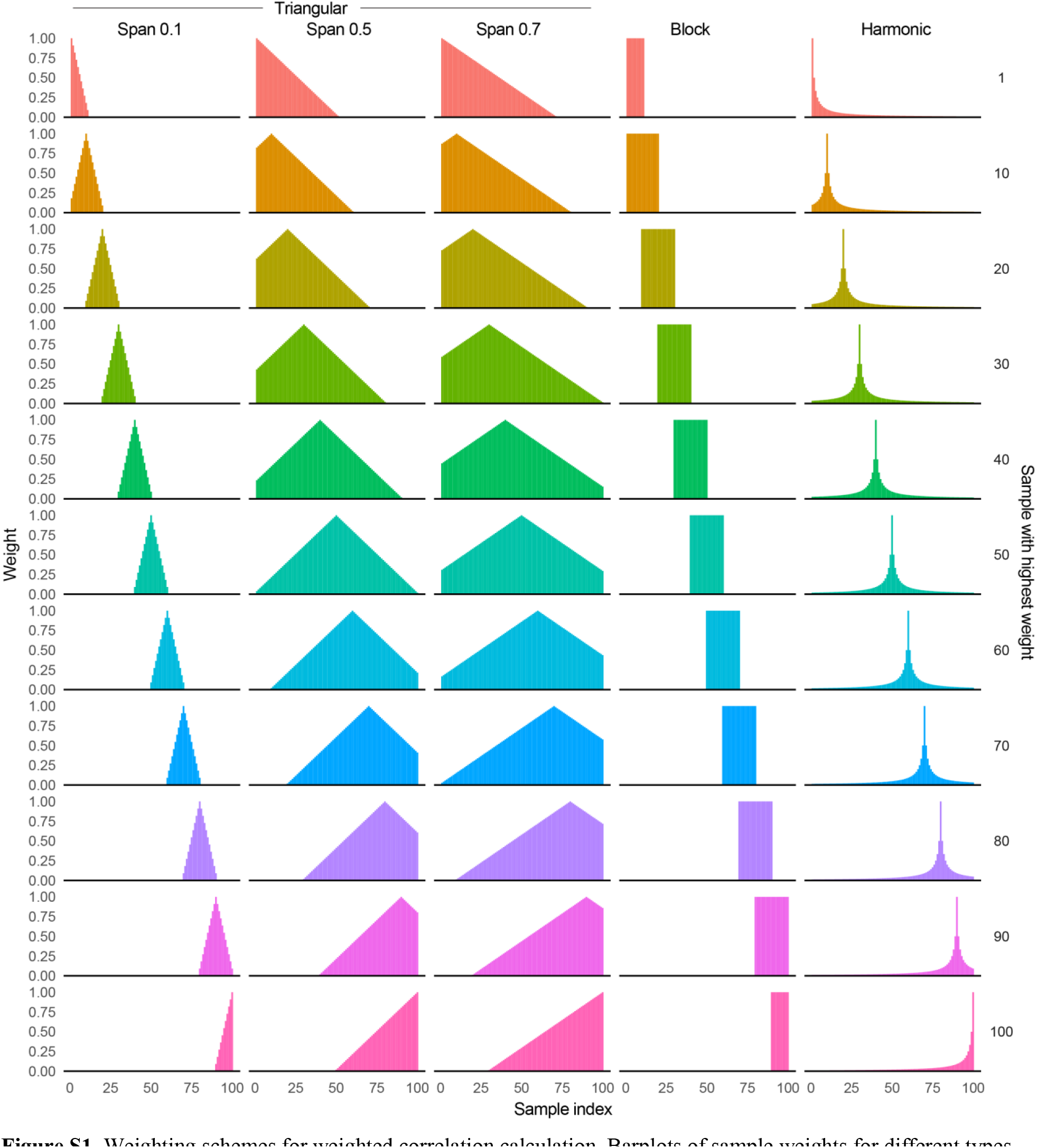
Weighting schemes for weighted correlation calculation. Barplots of sample weights for different types of weighting schemes, triangular, block and harmonic, for an example sample set of length 100, shown for a subset of weight vectors at 1, 10, 20, …, 100. Triangular and harmonic weighting schemes allow choice of span for the weights, shown at span 0.1, 0.5, and 0.7 for the triangular scheme and span of 0.1 for the block scheme.

**Figure S2.**
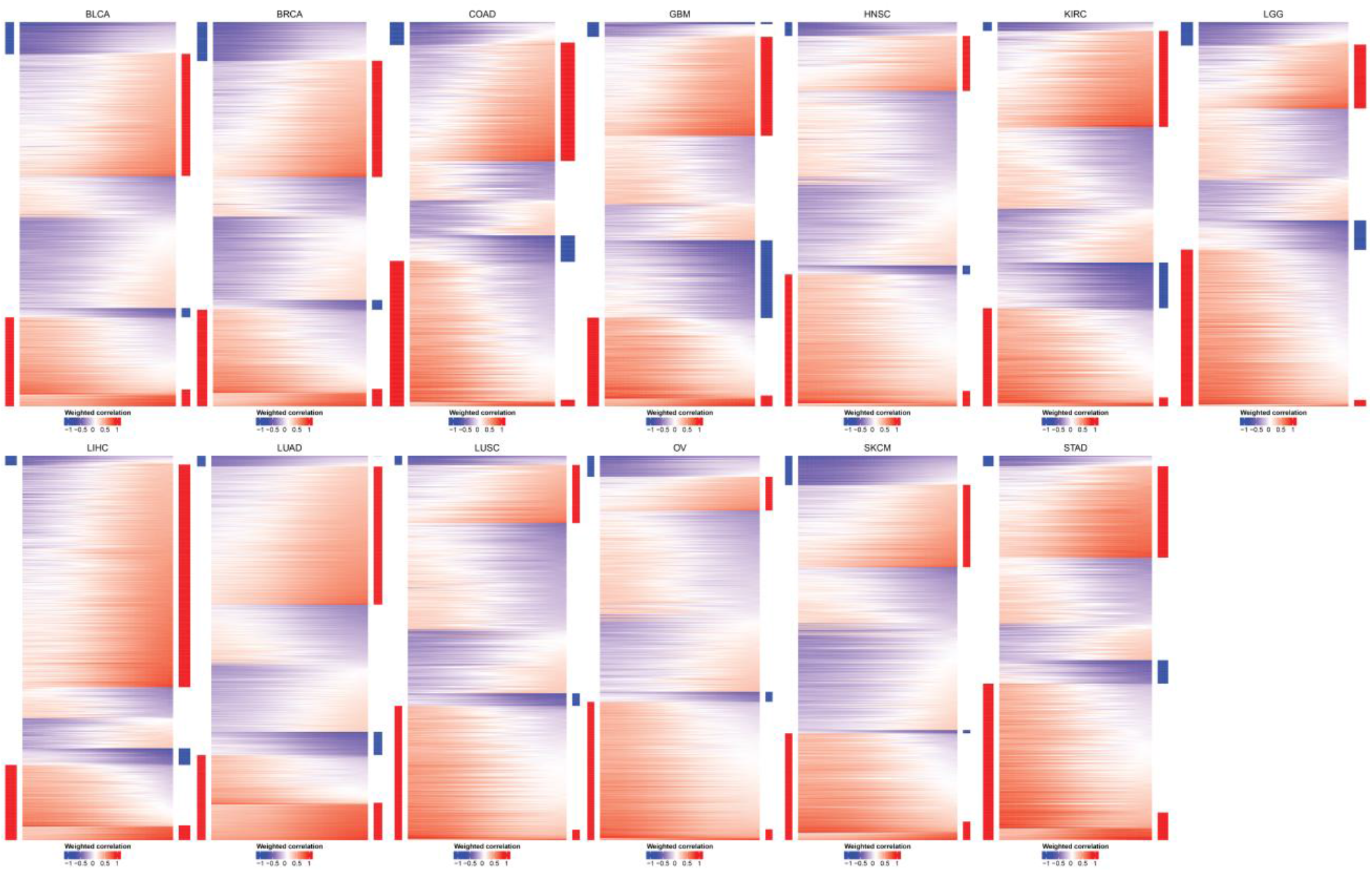
Heatmap of weighted correlations for significant gene pairs using DCARS across thirteen TCGA cancers. Heatmaps display the weighted correlation vectors from low to high survival samples (rows), with red indicating positive weighted correlation and blue negative weighted correlation, for all significantly DCARS gene pairs with non-zero change in weighted correlation. Coloured bars along the sides indicate gene pairs for which the weighted correlation is identified as positive (red) or negative (blue) or zero (white) at either low or high end of survival, thus characterising each gene pair in terms of the direction of change of weighted correlation Gene pairs identified as zero at both the low or high end of survival were removed.

**Figure S3.**
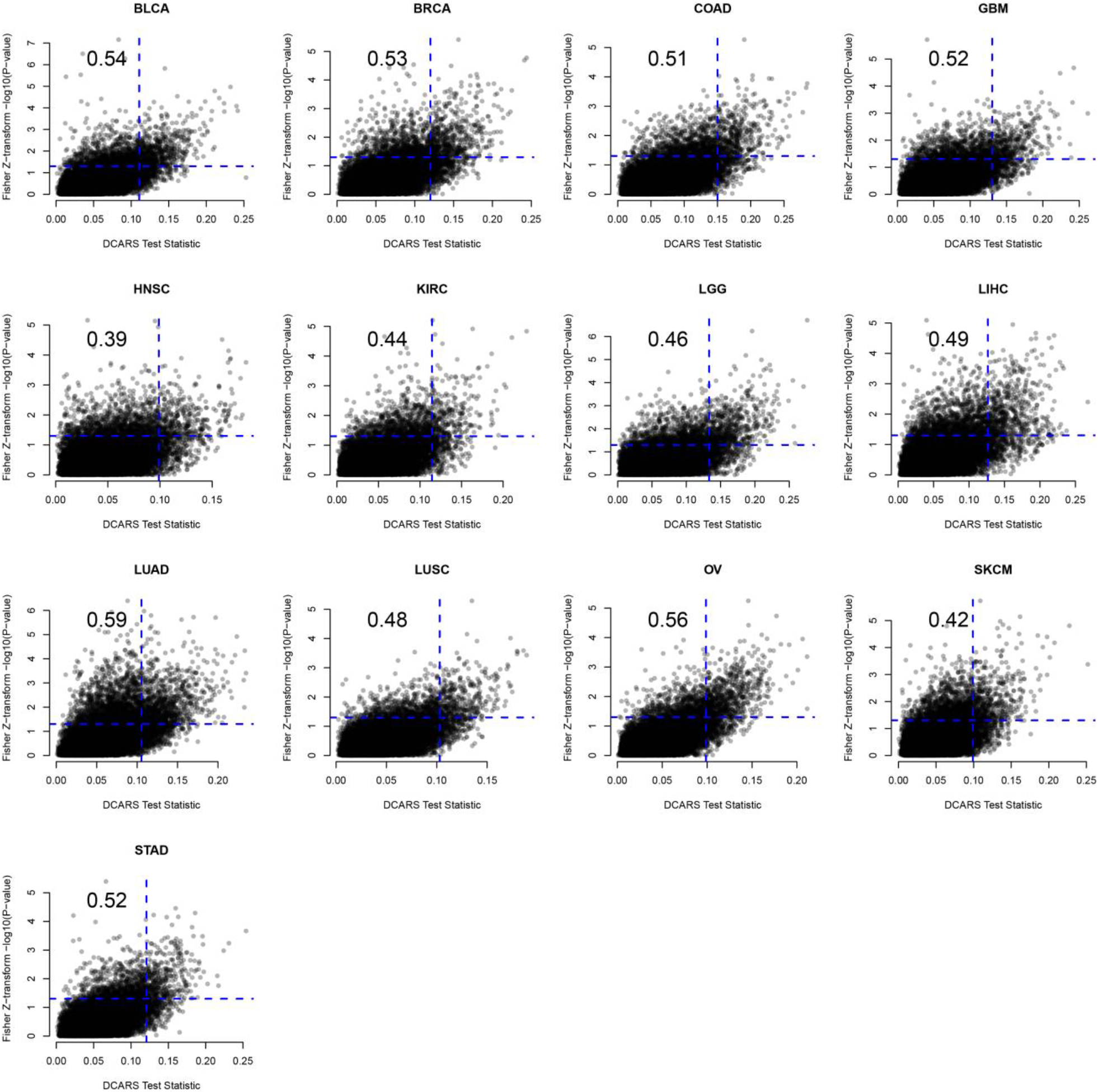
Scatterplots of DCARS test statistics vs −log(*p*-value) of Fisher’s Z-transformation test for 13 TCGA cancers. Spearman correlation is given in top right corner. Horizontal and vertical lines correspond to unadjusted *p*-value of 0.05.

**Figure S4.**
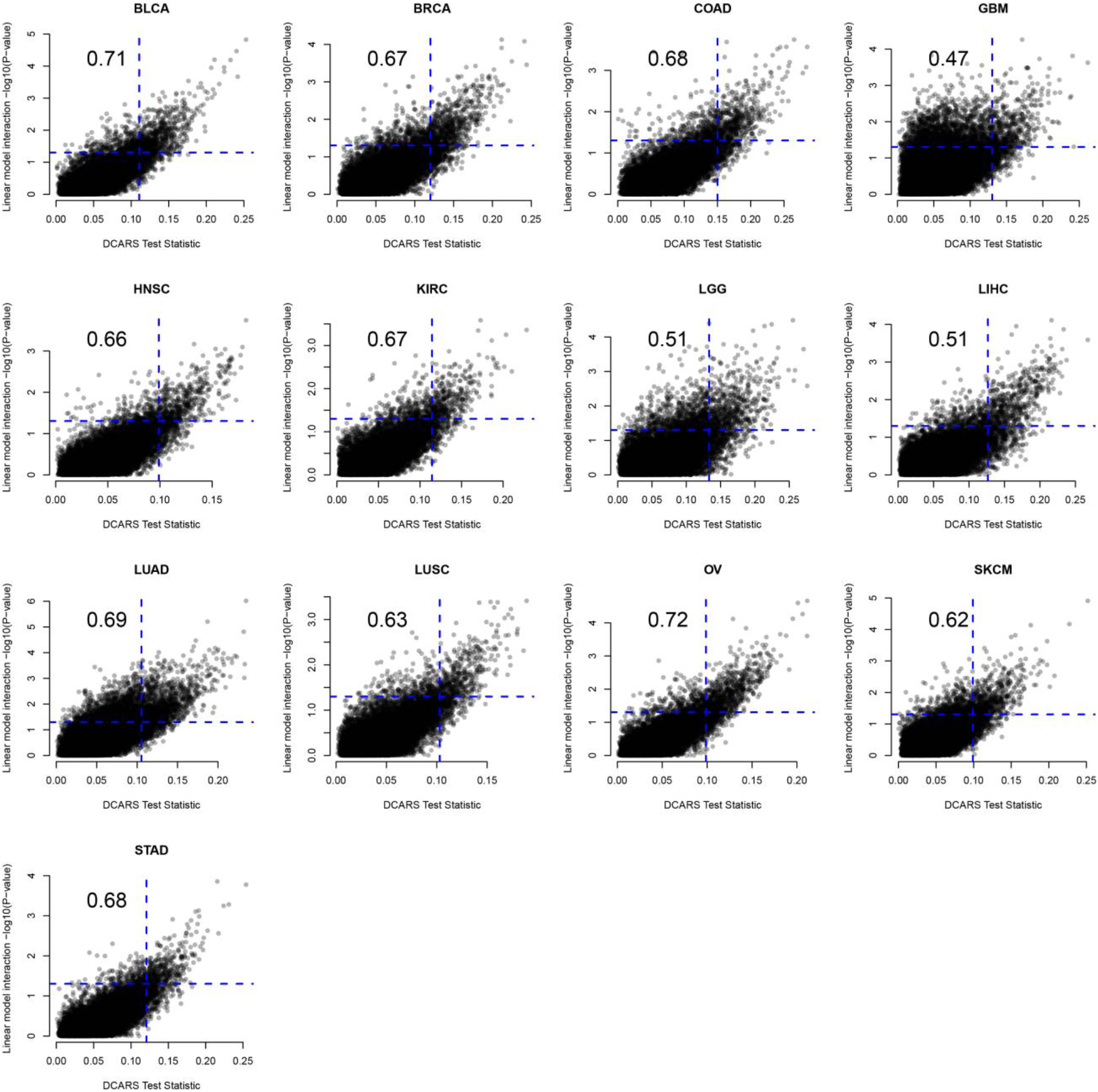
Scatterplots of DCARS test statistics vs −log(*p*-value) of Linear model interaction test for 13 TCGA cancers. Spearman correlation is given in top right corner. Horizontal and vertical lines correspond to unadjusted *p*-value of 0.05.

**Figure S5.**
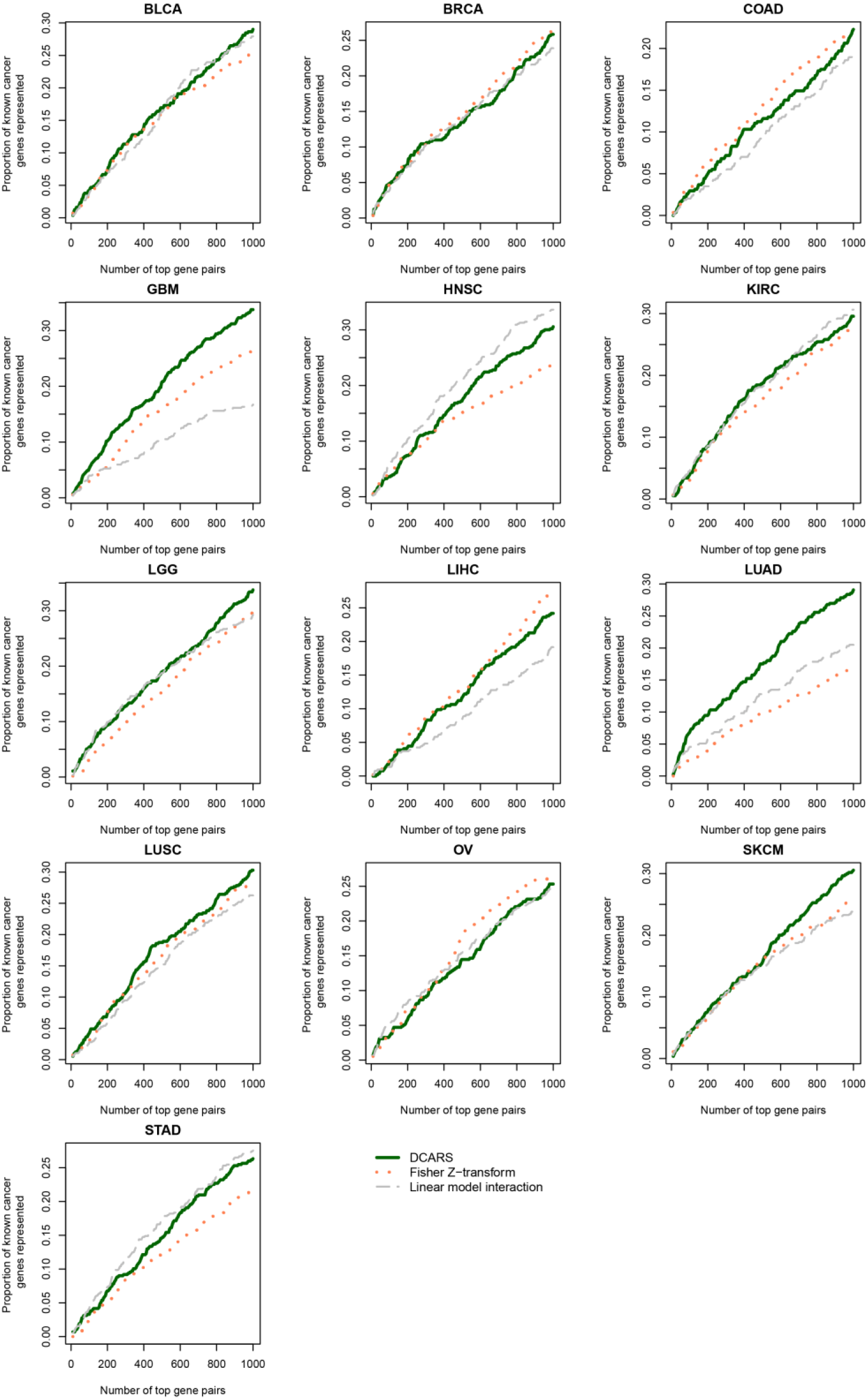
Line plots of the proportion of known cancer genes that appear in the top 1000 selected gene pairs using the DCARS (green solid), Fisher Z-transformation test (orange dotted) and linear model interaction test (grey dashed). A higher value indicates that a higher set of known cancer genes appear at least once in the top ranked list of gene pairs.

**Figure S6.**
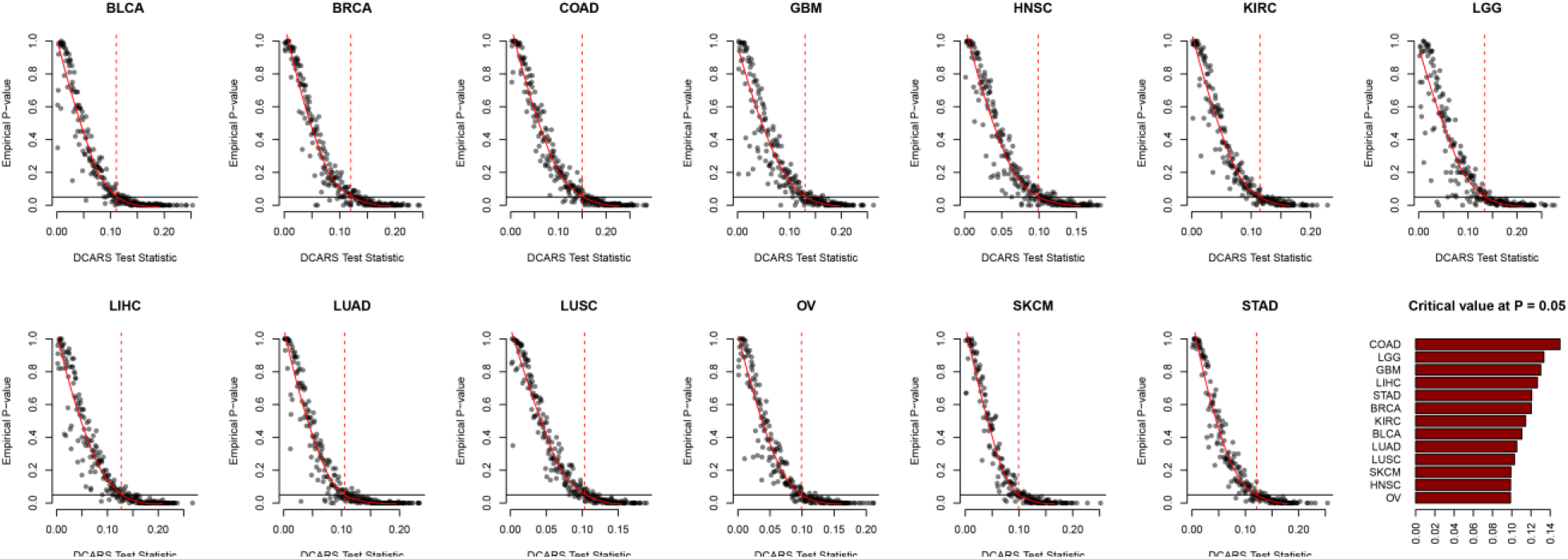
Observed test statistic and permutation based *p*-values for each cancer dataset. Once observed test statistics were calculated, a stratified sample of up to 500 gene pairs were taken to carry out permutation based testing with 100 permutations. A smoothed curve was fit for each cancer and the value associated with unadjusted *p*-value of 0.05 was taken to be the critical value for further interrogation, where gene pairs with a higher observed test statistic than the critical value were considered significantly differentially correlated across survival time.

**Figure S7.**
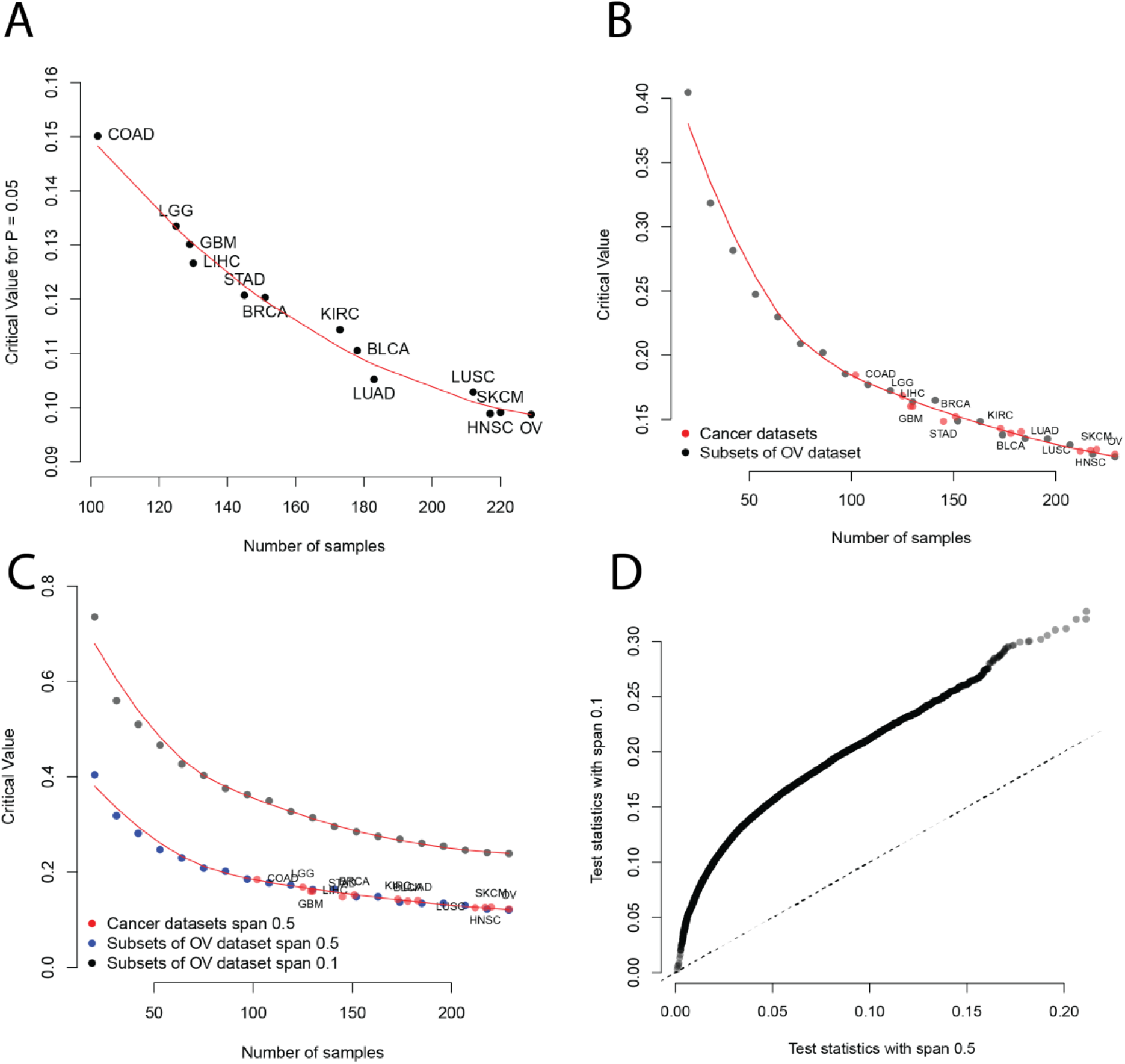
Comparison of critical test statistic values and number of samples. **A**. Number of samples and estimated test statistic critical value at P = 0.05 for the TCGA datasets. **B**. We took the TCGA dataset with the highest number of samples, OV, and successively took subsamples of this dataset, and calculated critical values as shown in Figure S1. This showed a monotonic trend of decrease in critical value as number of samples increased. Overlaying the graph from A and B shows the other cancers follow suit with the subsets of OV dataset. **C**. OV was subsampled and the DCARS test carried out with critical values found, but for a triangular weighting scheme with span of 0.1 (as opposed to 0.5), and the critical value appears to be parallel but lies above that of the 0.5 span. **D**. We graphed the observed test statistics using a triangular weighting scheme with span 0.1 and 0.5 for the OV dataset and find they are largely monotonically related.

**Figure S8.**
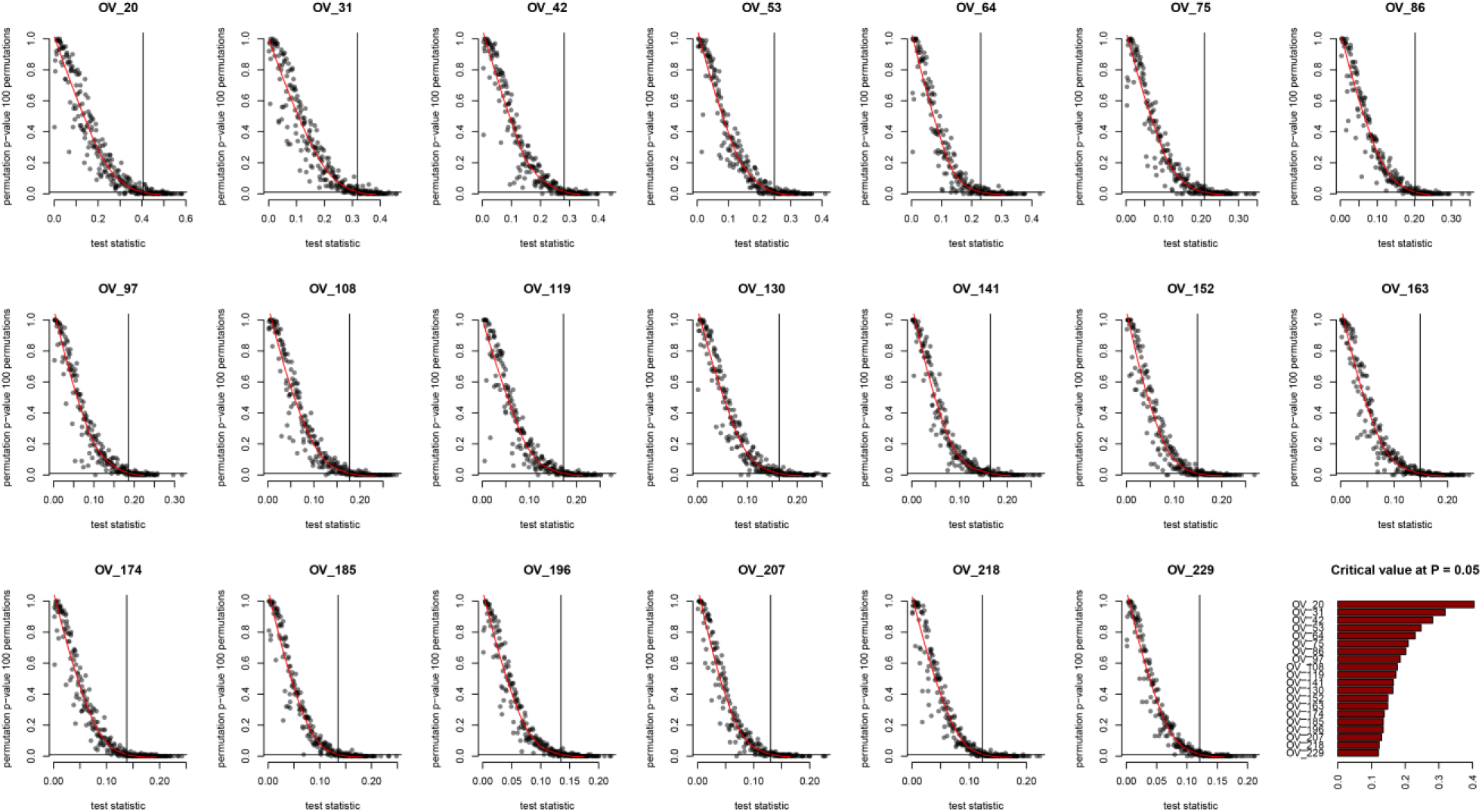
Observed test statistic and permutation based *p*-values for each subset of OV cancer dataset. For each subset of the 229 samples from the OV dataset, observed test statistics were calculated using a triangular weight matrix of span 0.5. Then, a stratified sample of up to 500 gene pairs were taken to carry out permutation based testing with 100 permutations. A smoothed curve was fit for each cancer and the value associated with unadjusted *p*-value of 0.05 was taken to be the critical value for further interrogation, where gene pairs with a higher observed test statistic than the critical value were considered significantly differentially correlated across survival time.

**Figure S9.**
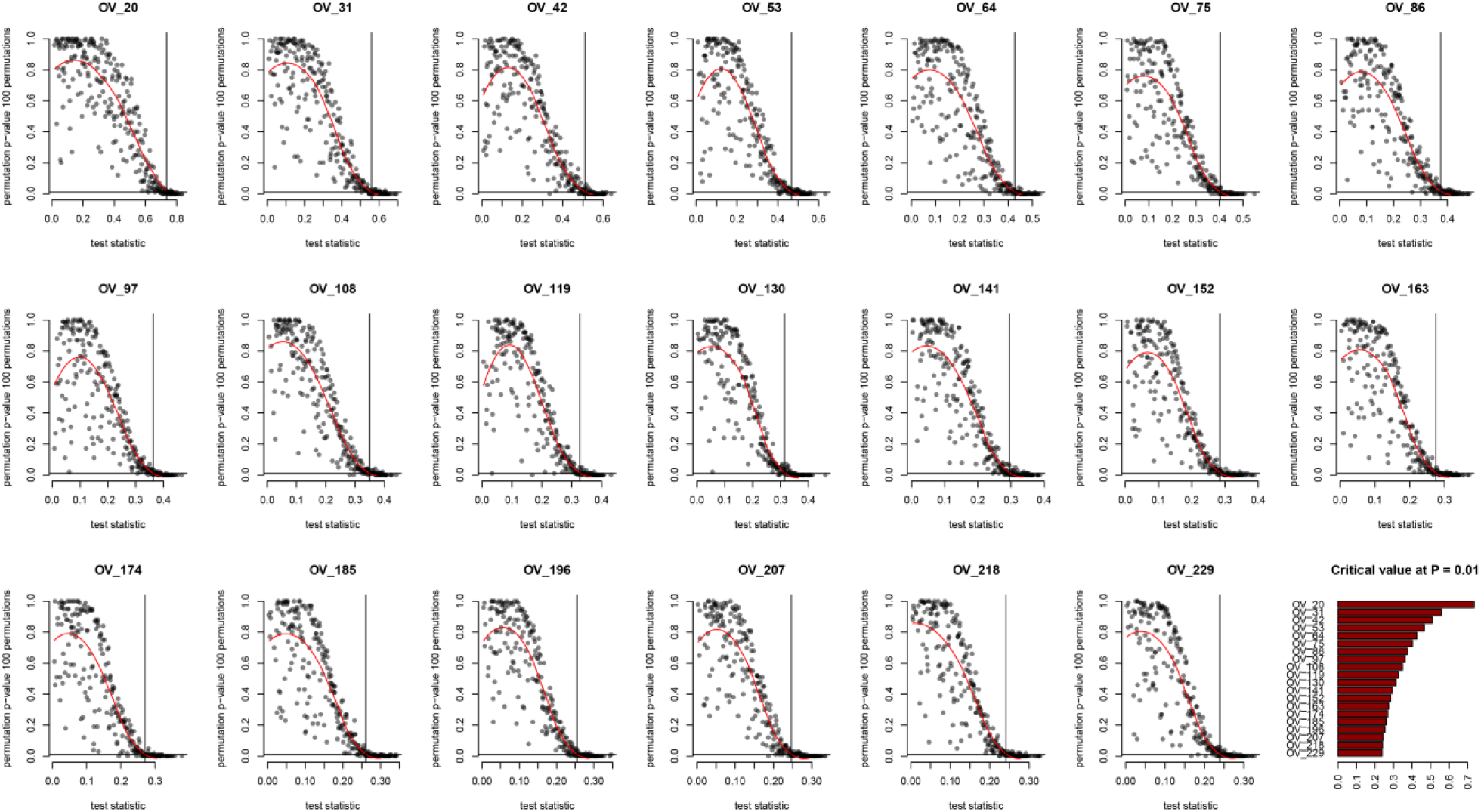
Observed test statistic and permutation based *p*-values for each subset of OV cancer dataset with triangular weight matrix with span 0.1. For each subset of the 229 samples from the OV dataset, observed test statistics were calculated using a triangular weight matrix of span 0.1. Then, a stratified sample of up to 500 gene pairs were taken to carry out permutation based testing with 100 permutations. A smoothed curve was fit for each cancer and the value associated with unadjusted *p*-value of 0.05 was taken to be the critical value for further interrogation, where gene pairs with a higher observed test statistic than the critical value were considered significantly differentially correlated across survival time.

**Figure S10.**
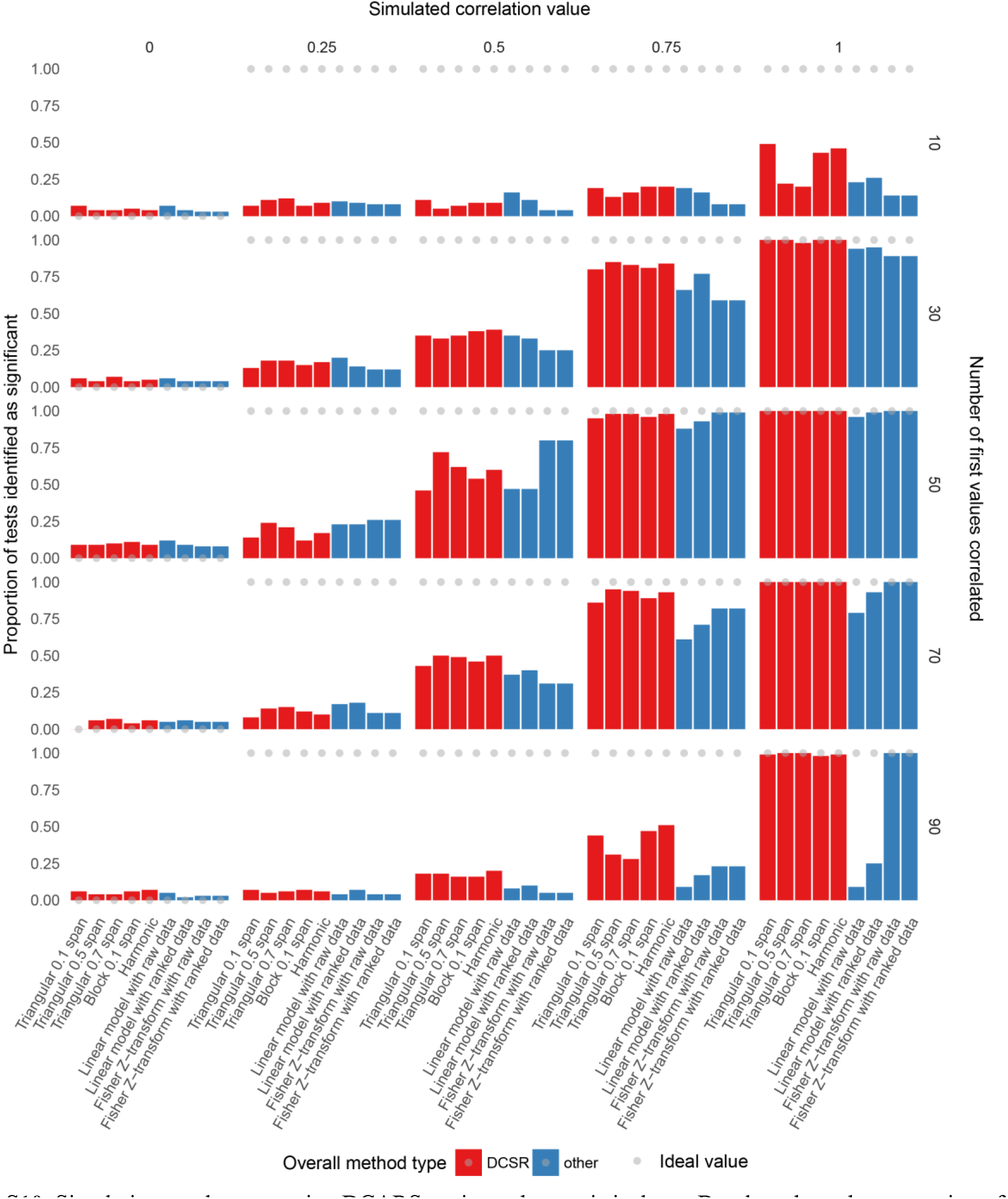
Simulation results comparing DCARS against other statistical test. Barplots show the proportion of tests for simulated data that were considered significant with a *p*-value of 0.05 across a set of 100 data points where the first set of 10, 30, 50, 70 or 90 values were correlated at values of 0, 0.25, 0.5, 0.75 and 1. Grey dots show the ideal values associated with a false positive rate of 0 and statistical power of 1. Simulations with correlated values of 0 adhered to the null model of no differential correlation across data points and thus estimate the false positive rate. DCARS methods perform similarly to other statistical tests in terms of estimated false positive rate. DCARS methods either perform similarly or outperform other statistical tests in other simulation scenarios, e.g. when the number of first values correlated is far from 50/100, and when simulated correlation values are 0.5 or less.

**Figure S11.**
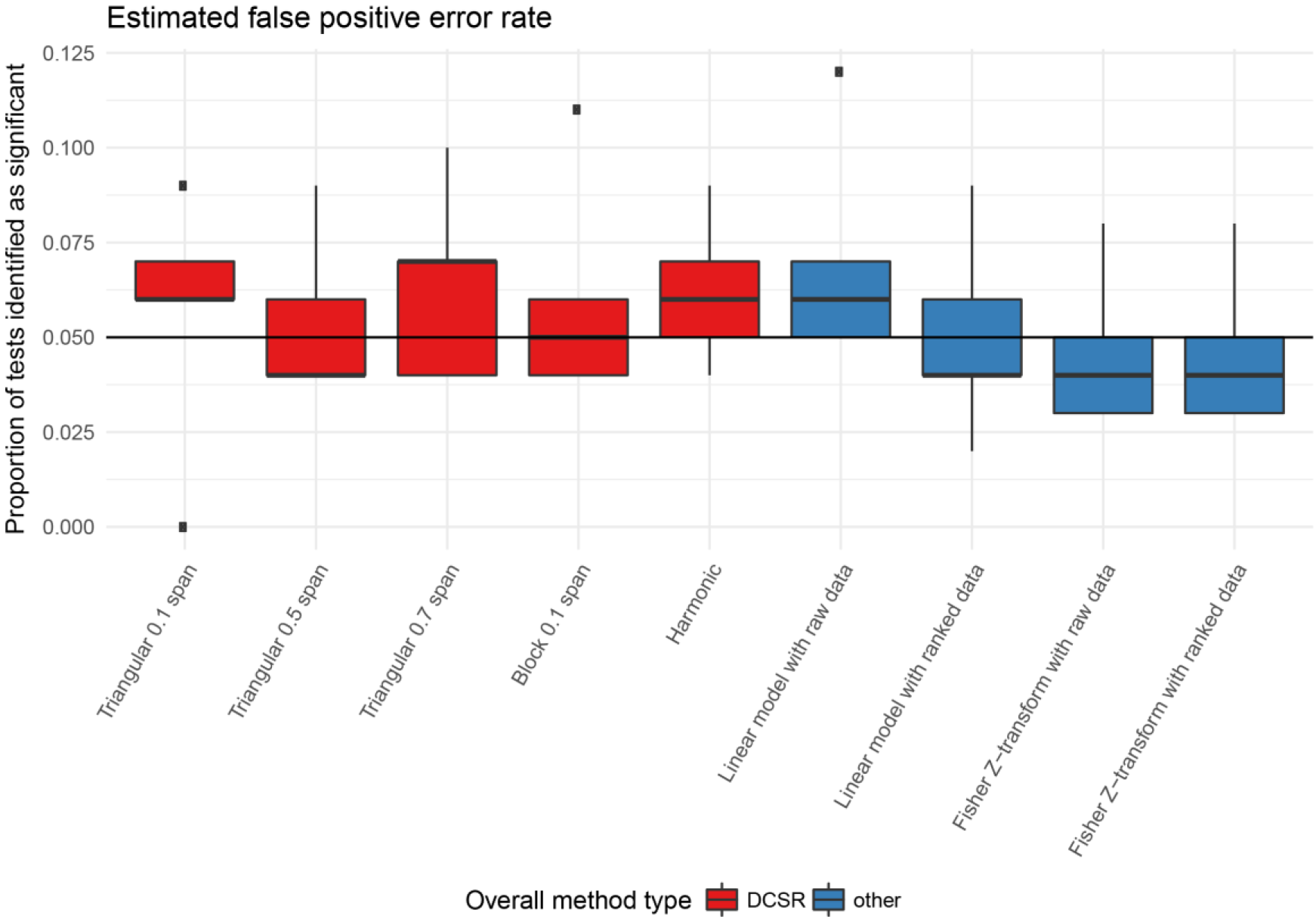
Boxplots of estimated false positive error rates in simulation study. Data were simulated to have no correlation and the proportion of ‘significant’ test results across 100 simulations are shown for DCARS methods and other statistical tests. All methods are similar in their estimated false positive rate.

**Figure S12.**
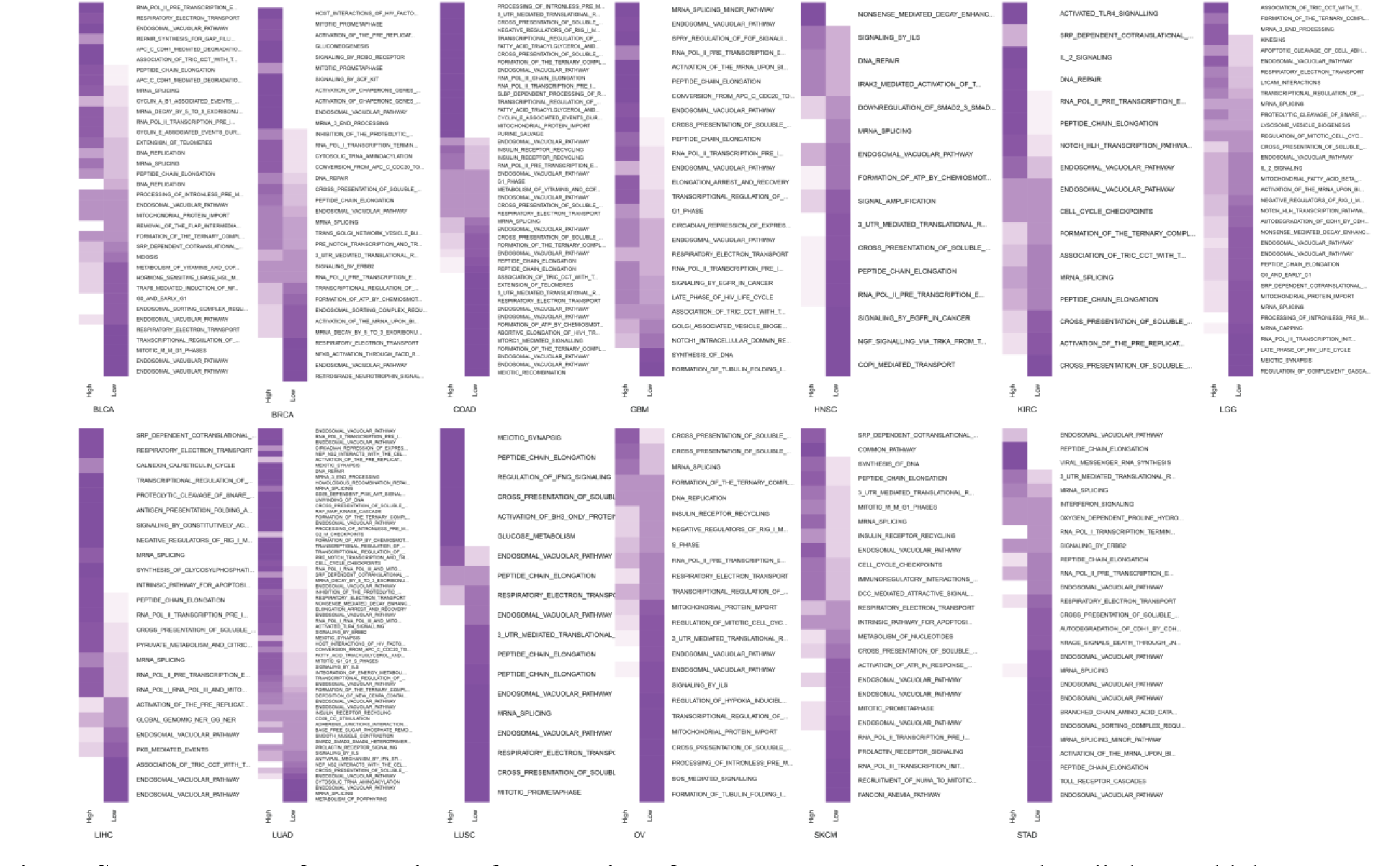
Heatmap of proportions of correlations for pathway per cancer. Purple cells have a higher proportion of gene pairs with correlation in either the high or low survival groups per subnetwork per cancer.

**Figure S13.**
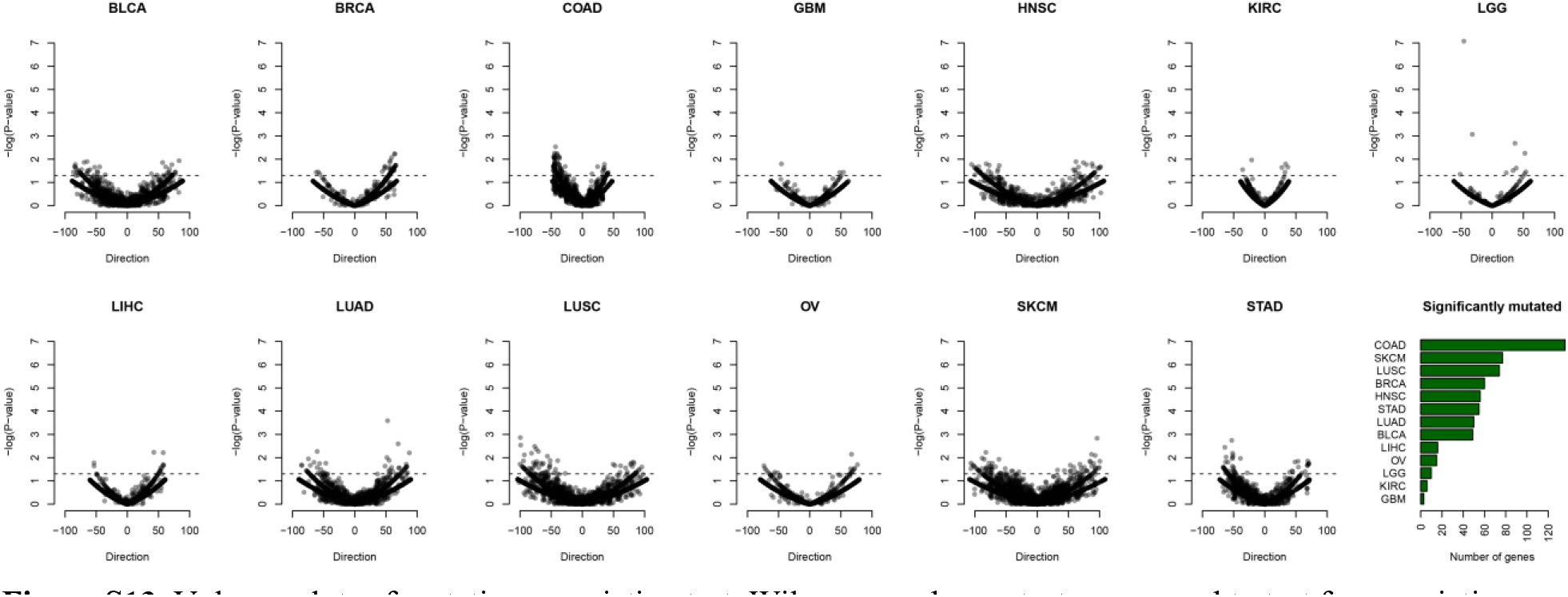
Volcano plots of mutation association test. Wilcoxon rank sum tests were used to test for association with survival ranking and presence of a non-silent mutation in the gene. Direction (x-axis) is given as the median survival ranking of samples without mutations in the gene subtracted by the median survival ranking of samples with mutations in the gene, thus higher values associate with genes with mutations in low survival samples. Y-axis represents −log10(*p*-value) of the two-sided Wilcoxon Rank Sum test.

**Supplementary files**

Supplementary files are located in https://github.com/shazanfar/DCARS.

**Supplementary File 1** – Zip file containing tab-separated text files for each tested gene pair across 13 cancer datasets, including the DCARS test statistic, Fisher Z-transform test, linear model interaction test as well as other information.

**Supplementary File 2** – Excel spreadsheet of gene pairs significant DCARS in two or more cancers.

**Supplementary File 3** – Excel spreadsheet of input gene list and output Biological Process Gene Ontology Enrichment Analysis performed by STRING database (v 10.5).

**Supplementary File 4** – PDF file of network diagrams of DCARS gene pairs for all 13 TCGA datasets.

